# Myocarditis and neutrophil-mediated vascular leakage but not cytokine storm associated with fatal murine leptospirosis

**DOI:** 10.1101/2024.10.01.616081

**Authors:** Stylianos Papadopoulos, David Hardy, Frédérique Vernel-Pauillac, Magali Tichit, Ivo G. Boneca, Catherine Werts

## Abstract

Leptospirosis is a neglected re-emerging zoonosis caused by *Leptospira* spirochetes. Its pathophysiology remains mysterious, especially in the case of severe infection with *L. interrogans*.

In the field of infectious diseases, the cause of death is rarely investigated in preclinical models. Here, for the first time, we identified unanticipated organ failures associated with death in a murine model of acute leptospirosis.

Despite clinical similarities between bacterial sepsis and leptospirosis, striking differences were observed. Neither lung, liver, or kidney injury nor cytokine storm, or massive necroptosis could explain death. In contrast, severe leptospirosis was associated with high serum levels of the anti-inflammatory cytokine IL-10 and the chemokine RANTES, neutrophilia, pancreatitis and vascular damage. Unexpectedly, we demonstrated neutrophil-induced vascular permeability, making neutrophils a potential new therapeutic target. Strikingly, the main cause of death was myocarditis, an overlooked complication of human leptospirosis.

These features are also found in patients, making this model a paradigm for better understanding human leptospirosis and designing novel therapeutic strategies.

## Introduction

Leptospirosis is a largely neglected zoonosis caused by the spirochete bacteria belonging to the genus *Leptospira* that severely affects more than 1 million people and is responsible for nearly 60,000 deaths per year worldwide (1). Pathogenic leptospires, with *Leptospira interrogans* (*L. i*) being the most virulent member of the family, are usually transmitted by chronically infected animals that shed the bacteria in their urine and contaminate the environment, mostly water and soil. Due to climate change, the prevalence of this disease is currently increasing, especially in tropical countries affected by floodings, which favors the spread and infection of leptospires (2). Indeed, pathogenic leptospires penetrate the skin and mucosa and disseminate in the circulation. They are resistant to complement (3) and can replicate in the blood and affect all organs, particularly the liver, lungs, kidneys and brain (4). Leptospirosis is common in East Asia and Americas among rice field and sugar cane workers, and in large urban areas where sanitation is poor and proximity to reservoir animals is favored (1). There are more than 60 species of leptospires and 300 of serovars (5), making vaccine development challenging. Besides, the broad spectrum and diversity of clinical manifestations makes diagnosis of leptospirosis difficult. Indeed, once infected, humans may develop either an asymptomatic or mild flu-like illness with symptoms common to many other infectious diseases (fever, chills, myalgias, nausea), but in some cases, humans may be susceptible to a potentially lethal form of leptospirosis characterized by jaundice, acute respiratory distress syndrome (ARDS), septic shock, and multi-organ dysfunction syndrome (MODS)(6).

The exact reason for the death of most patients with leptospirosis is still unclear. Different reviews mention the involvement of a potential cytokine storm in the acute phase of leptospirosis, leading to hyperinflammation and ultimately sepsis (4, 7). Sepsis is defined as a “life-threatening dysfunction caused by a dysregulated host response to infection” (8), and a plethora of molecular players, cellular types and compartments are implicated in its pathogenesis. Upon infection, the host innate immune system first recognizes microbial-associated molecular patterns (MAMPs) that bind to pattern recognition receptors (PRRs), such as Toll-like receptors (TLRs). This MAMP/PRR recognition triggers an inflammatory response with the secretion of cytokines and chemokines involved in the recruitment of leukocytes to the site of infection. In the best case, the pathogen is rapidly cleared by phagocytes, but an exaggerated and uncontrolled immune response can lead to a cytokine storm. The main players in this deleterious outcome are TNF, IL-1β and IFN-γ, all of which can activate cells by binding to their specific receptors, thereby amplifying inflammation and producing more cytokines, resulting in the so-called cytokine storm (9). Nevertheless, in the case of *Leptospira* this innate immune recognition is poor as these bacteria have managed to develop unique strategies to escape the sensing of many PRR pathways of the innate host defense (10). Apart from the activation of TLR2 by the leptospiral lipoproteins, the lipopolysaccharide (LPS) escapes the activation of human TLR4 and it only partially activates the murine TLR4 (11). Other mechanisms linked to LPS also dampen the IL-1β release and inflammatory and immune responses (12–14). Furthermore, the intracellular localization of the leptospiral endoflagella impairs its sensing by TLR5 (15) and the tight binding of LipL21, one of the major leptospiral lipoproteins, to peptidoglycan prevents its recognition by the NOD receptors (16).

We have previously shown that sublethal infection with 2×10^7^ *L. i* Manilae L495 per mouse (or its bioluminescent derivative strain MFLum1) causes a biphasic disease characterized by a 7-day acute but self-resolving phase of blood dissemination of the bacteria, followed by their chronic, stable and asymptomatic renal colonization in the proximal tubules. However, the use of the higher dose of 10^8^ MFLum1 leads to the death of mice 3 to 4 days after infection (17). We have also shown that *L. i* Icterohaemorrhagiae Verdun does not colonize mouse kidneys and is only responsible for non-lethal infections, although it may cause severe weight loss and jaundice, at the higher dose of 5×10^8^ bacteria (18). To investigate the pathophysiological and inflammatory responses during acute leptospirosis in mice, we compared infection with these two serovars to highlight differences that may provide insight into the causes of lethality.

## Materials and Methods

### Leptospira interrogans culture

The pathogenic *Leptospira interrogans* (*L. i*) serovar Manilae strain L495 and the serovar Icterohaemorrhagiae strain Verdun (Virulent cl3) (18), both originally obtained from human patients with leptospirosis in the Philippines and France, respectively, were grown in liquid Ellinghausen-McCullough-Johnson-Harris (EMJH) medium at 30°C without agitation. They were passaged weekly to a final concentration of 10^6^ bacteria/mL to be used one week later at the end of the exponential phase. For infection, leptospires were centrifuged at 3,200 x *g* for 25 min at room temperature, resuspended in endotoxin-free PBS (Lonza), and counted using a Petroff-Hauser chamber. For *in vivo* infection of mice, *L. i* between passage 3 and 12 were used.

### *In vivo* infection of mice

Adult (6-10 weeks old) C57BL/6J mice were obtained from Janvier Labs and maintained under standard conditions in the animal facility of the Institut Pasteur for at least 1 week prior to infection. For each infection, groups of 3 or 5 mice (both male and female) were infected intraperitoneally (IP) with 10^8^ *Leptospira*/mouse in a final volume of 200 μL, diluted in PBS. Control groups of uninfected naive animals were injected with 200 μL PBS. Body weight changes, temperature and clinical signs were monitored daily. The clinical score was determined as a composite of weight loss, alertness, locomotion, and body function as described (18). On day 3 post-infection (p.i.), mice were euthanized by cervical dislocation or progressive CO_2_ inhalation. Blood, urine, and organs were snap frozen in liquid nitrogen and stored at −80 °C or used fresh, depending on the experimental design. For serum or plasma collection, blood samples were collected by cardiac puncture into serum separation tubes (Microvette 200 serum-gel, Sarstedt TPP, Fisher Scientific) or tubes containing 20µl of EDTA 100 μM, and after centrifugation, serum/plasma was separated, aliquoted, and stored at −20 °C. Urine was collected just prior to animal sacrifice. For the injection of heat-killed (HK) bacteria, both *Leptospira* serovars and *Escherichia coli* K12 were centrifuged at 3,200 x *g* for 30 min at room temperature, resuspended in PBS, counted in a Petroff-Hauser chamber and finally heat-inactivated at 65 °C for 30 min. The mixture of HK bacteria-PBS or PBS alone was allowed to cool at room temperature and injected into the mice by the IP route at 10^8^ bacteria/mouse. Clinical signs were observed every 30 minutes up to 4 hours after injection. Mice were euthanized by cervical dislocation and blood was collected in tubes containing EDTA (100 μM). Plasma was collected after centrifugation and stored at −20 °C for further analysis.

### Ethic statement

All experiments were performed in accordance with the guidelines established by the French and European regulations for the care and use of laboratory animals and approved by the Institut Pasteur Committee on Animal Welfare (CETEA) protocol dap210024 /APAFIS#30877-2021040212335074 v1.

### Morphological and immunohistochemical analysis

Organs (liver, lung, spleen, pancreas, brain and intestine) were harvested on day 3 p.i. and directly fixed in 10% neutral buffered formalin for 48 hours or kidneys for 2 hours in Dubosq-Brazil, followed by post-fixation in 10% formol (19). Fixed tissues were further washed in 70% ethanol and embedded in paraffin. Tissue sections were stained with hematoxylin-eosin (H&E) to evaluate inflammatory changes by light microscopy. Immunohistochemical studies were performed using antibodies against macrophages, neutrophils, T lymphocytes, caspase-3 and *Leptospira* (Table 1). Stained slides were evaluated with ZEISS AxioScan.Z1 Digital Slide Scanner and Zen software (Zeiss).

### Leptospiral loads determination with qPCR

Quantification of leptospiral DNA was performed as previously described (18). Total leptospiral DNA was extracted from blood and organs (liver, kidney, lung, spleen, pancreas and brain) collected at day 3 p.i. using the QIAamp DNA Kit (Qiagen). Briefly, organs were homogenized at room temperature in a mechanical disruption apparatus (Retsch MM400) using 2.8 mm diameter metal beads. The concentration of purified DNA was quantified using Nanodrop. Leptospiral DNA was specifically targeted using primers and probes designed in the *lpxA* gene conserved in *Leptospira interrogans* serovars (Table 2). Normalization was performed using the *nidogen* gene (Table 2). qPCR was performed on a StepOne Plus real-time PCR machine (Applied Biosystems) using the following settings (absolute quantification program) with the following conditions for FAM TAMRA probes 50 °C for 2 min, followed by 95 °C for 10 min and 40 cycles of 95 °C for 15 s and 60 °C for 1 min. Results are expressed as the number of leptospires per 50 μL of blood or per 200 ng of total DNA extracted from organs. The limit of detection is 100 leptospires/200 ng DNA.

### Determination of cytokine mRNA expression by RT-qPCR

Total RNA was extracted from organs (liver, kidney, lung, spleen, heart) harvested on day 3 p.i. using the RNeasy Mini Kit (Qiagen). Briefly, organs were homogenized at 4 °C using Bead Tubes type F (Macherey-Nagel) and the lysate was further used for RNA isolation. The concentration of purified RNA was quantified using Nanodrop. One µg of RNA was subjected to reverse transcription using Superscript II reverse transcriptase (Invitrogen) according to the manufacturer’s recommendations. Generated cDNAs were either stored at −20°C or used for qPCR on a StepOne Plus real-time PCR machine (Applied Biosystems) with primers and probes targeting murine hypoxanthine-guanine phosphoribosyltransferase (*hprt)*, used as a housekeeping gene, *il1b, il6, il10, ifng, inos, tnf,* and *rantes* (Table 2). The settings used (relative quantification program) with the following conditions for FAM-TAMRA probes were 50 °C for 2 min, followed by 95 °C for 10 min and 40 cycles of 95 °C for 15 s and 60 °C for 1 min. The results were analyzed using the comparative 2^-ΔΔCt^ method with a first normalization by the internal *hprt* control and a second normalization by the PBS control group.

### Cytokine and biochemical markers determination with ELISA

For cytokine determination, serum/plasma and organs (liver, kidney, lung, spleen, pancreas, and heart) collected on day 3 p.i. were weighed and lysed in a buffer containing 200 mM NaCl, 10 mM Tris-HCl, 5 mM EDTA, 10% glycerol, 1 mM PMSF, with complete anti-protease mixture tablets (Roche) to a final concentration of 100 mg/mL. After mechanical disruption in a Retsch MM400 apparatus using 2.8 mm diameter metal beads at 4 °C and centrifugation at 16,000 x *g* for 10 min at 4 °C, the lysates were stored at −80 °C until cytokine dosing. The concentration of TNF, IL-1β, IL-6, IFN-γ, IL-10 and RANTES in the supernatants was determined using ELISA kits (R&D Systems) according to the manufacturer’s recommendations. Sera collected on day 3 p.i. were used for the measurement of biochemical markers. Lipocalin-2/neutrophil gelatinase-associated lipocalin (NGAL, renal marker, R&D Systems), cardiac troponin I (cTnI, cardiac marker, Abcam), creatinine kinase-myoglobin binding (CK-MB, cardiac marker, Abcam), and aldosterone (dehydration marker, Abcam) were quantified by ELISA according to the manufacturer’s recommendations.

### Determination of biochemical parameters

Serum or urine collected on day 3 p.i., either fresh or stored at −20 °C, were used for the determination of vital biochemical parameters of mouse organ function. For liver function, serum total bilirubin (MAK126, Sigma), alanine aminotransferase activity (ALAT, MAK052, Sigma) and aspartate aminotransferase activity (ASAT, MAK055, Sigma) were quantified. For renal function, serum creatinine (MAK080, Sigma) was measured. For pancreatic function, lipase activity (MAK046, Sigma) and amylase activity (MAK009, Sigma) were measured in serum and, for the latter, in urine. All colorimetric assays were performed according to the manufacturer’s instructions. Lactate dehydrogenase (LDH) release was quantified in serum and organ lysates (liver, kidney, lung, spleen, pancreas, and heart) collected on day 3 p.i., diluted to a final concentration of 500 μg/mL, using CyQuant LDH Fluorometric Assays (Invitrogen) according to the manufacturer’s instructions. Fluorimetry was measured at 560 nm (excitation) / 590 nm (emission) on a TECAN Spark fluorimeter (Life Sciences).

### Flow cytometry

Blood and organs (liver, kidney, lung, spleen) collected at day 3 p.i. were directly homogenized and stained for flow cytometry analysis. Briefly, blood was collected in tubes containing 20µl of EDTA 100 μM and red blood cells were lysed using 1x red blood cell lysis buffer (Biolegend) for 5 min at room temperature. After centrifugation at 400 x *g* for 7 min, cells were washed in PBS supplemented with 0.5% fetal bovine serum (FBS) + 0.4% EDTA 0.05 M. Single cell suspensions from blood were counted and prepared for surface staining in 96-well plates (U-bottom, TPP). For organ analysis, freshly harvested tissues were incubated with appropriate enzymes (Miltenyi Biotec, Table 3) followed by mechanical dissociation (gentleMACS, Miltenyi Biotec). Homogenates were passed through 100 μm strainers (Miltenyi Biotec) in PBS supplemented with 0.5% FBS + 0.4% EDTA 0.05 M, then lysed in red blood cell lysis buffer (Sigma) for 5 minutes at room temperature and successively passed through 70 μm and 30 μm strainers (Miltenyi Biotec). For kidney samples, an additional enrichment step was performed by density gradient centrifugation. Specifically, total cell isolates were resuspended in 4 mL of 70% Percoll solution (GE Healthcare), overlaid with 4 mL of 37% Percoll, followed by 1 mL of 30% Percoll. The gradient was centrifuged at 900 x *g* for 30 min at room temperature and the leukocyte enrichment was collected at the interface of the 70%-37% layers. For all, cells were incubated for 30 min at 4 °C with anti-mouse CD16/CD32 to block non-specific binding, followed by incubation for 30 min at 4 °C in a cocktail of surface staining labeling antibodies in PBS supplemented with 0.5% FBS + 0.4% EDTA 0.05 M (Table 4). Cells were then stained for viability with the fixable viability dye eFluor780 (65-0865-14, Invitrogen) for 5 minutes at 4 °C and then fixed with 4% w/v paraformaldehyde in PBS for 5 minutes at room temperature. After a final wash in PBS supplemented with 0.5% FBS + 0.4% EDTA 0.05 M, samples were acquired on an LSR Fortessa (BD Biosciences) flow cytometer and data analysis was performed using FlowJo software v10 (TreeStar). The gating strategy used is adapted from (20).

### *In vivo* neutrophil depletion

Neutrophils were depleted in C57BL/6J mice by retro-orbital injection of 100 µL of 50 μg anti-mouse Ly6G/Ly6C (GR-1) monoclonal antibody clone RB6-8C5 (16-5931-85, eBioscience) 24 h before infection. Control animals were injected with the appropriate rat IgG2b κ isotype control (16-4031-85, eBioscience) under the same conditions. Prior to neutralizing antibody injection, mice were anesthetized with a constant flow of 2.5% isoflurane mixed with oxygen and air according to the manufacturer’s recommendations using an XGI-8 anesthesia induction chamber (Xenogen Corp.). Mice were maintained in the anesthesia chamber for at least 10 minutes to allow adequate distribution of the injected substrate. Neutrophil depletion was confirmed by flow cytometric analysis of blood (see Table 4 for antibodies used for phenotyping).

### Evaluation of vascular permeability with Evans blue

On day 3 p.i., anesthetized (see previous paragraph for anesthesia details) mice (neutrophil-depleted or not) received an intravenous injection of 100 μL sterile 0.5% w/v Evans blue dye in PBS via the retro-orbital route, which was allowed to circulate for 30 min. Mice were euthanized by progressive inhalation of CO_2_ for 4 minutes and then perfused with 20 mL heparin (2 units/mL) in PBS. Organs (liver, kidney, lung, spleen) were then collected, weighted, and placed in 2 mL tubes containing 500 μL formamide. The samples were incubated at 37 °C for at least 48 hours to extract Evans blue dye from the tissues and centrifuged to pellet any remaining tissue fragments. The optical density at 620 nm of the supernatants was measured using a spectrophotometer, and the Evans blue concentration (ng/mL) of each sample was determined.

### Blood culture

Total blood (100 μL) was collected in tubes containing 20 µl of EDTA 100 μM by cardiac puncture at day 3 p.i. from mice (neutrophil depleted or not) and transferred directly to melted agar containing Brain Heart Infusion (BHI) medium tubes. After thorough homogenization, the blood/BHI mixture was poured into special long glass tubes that allow the growth of both anaerobic (bottom) and aerobic (top) bacteria. The tubes were incubated at 37 °C for at least 2 weeks to test for bacterial growth. Blood mixed with feces was used as a positive control of bacterial colony formation.

### Statistics

All graphs and statistical analyses were performed using GraphPad Prism 10 (GraphPad Software). No statistical method was used to power the sample. Comparisons between two groups were made using unpaired, two-tailed T-tests. 1-way ANOVA or 2-way ANOVA was used to compare more than two groups. Tests were considered significant at a *p*-value < 0.05 (\**p* < 0.05; \*\**p* < 0.01; \*\*\**p* < 0.001).

## Results

### *L. i* Manilae L495, but not *L. i* Ictero Verdun, causes mouse death

To compare the outcome of leptospirosis induced by two serovars of *Leptospira interrogans* (*L. i*), we intraperitoneally (IP) infected both male and female adult C57BL/6J mice with 10^8^ *L. i* Manilae L495 and *L.i* Icterohaemorrhagiae Verdun and followed and scored their symptoms over time according to a previously established clinical chart (18). As expected, individuals injected with the strain L495 reached the ethically humane endpoint at day 3 p.i. as they developed signs of acute leptospirosis, with lethargy, prostration, icteric urine, piloerection, and small eyes being the main clinical manifestations of severe lethal leptospirosis (Fig. 1A). However, at the same infection dose, mice in the Verdun group developed only mild symptoms with slight hematuria (Fig. 1A). This was confirmed by measurements of body weight and temperature changes during infection. Indeed, at day 3 p.i., lethally infected L495 mice lost on average 12.2% (females) and 15.0% (males) of their initial body weight compared to the uninfected PBS group and their temperature dropped from approximately 37.5°C to less than 32°C, whereas Verdun-infected mice lost on average only 1.9% (females) and 4.7% (males) of their initial body weight and their temperature recovered (Fig. 1B). Interestingly, we observed a slight sex difference between female and male mice in the Manilae group, with the latter being more susceptible to acute leptospirosis (Fig. 1A, 1B). In addition, we measured some characteristic markers of leptospirosis at day 3 p.i., such as serum bilirubin and urinary amylase, which may indicate liver and kidney dysfunction, respectively, as frequently observed in patients with leptospirosis (21). The results show that these markers were indeed elevated in mice infected with the L495 strain, but not with Verdun (Fig. 1C). We also measured serum release of the enzyme lactate dehydrogenase (LDH) a marker of cell death associated with inflammatory diseases (12). Compared to the control group, LDH was elevated in both Manilae and Verdun-infected mice, with a higher release in the Manilae group (Fig. 1C). Because the disease in Manilae infected mice was characterized by dramatic and very rapid loss in body weight and temperature, and because urine collection was extremely difficult, we speculated that dehydration might also contribute to the lethal phenotype. We measured serum aldosterone, a hormone involved in renal homeostasis and function, and a marker of hypokalemia often observed in patients with leptospirosis (22). Elevated aldosterone levels were measured only in the Manilae infected group (Fig. 1C).

**Figure 1.**
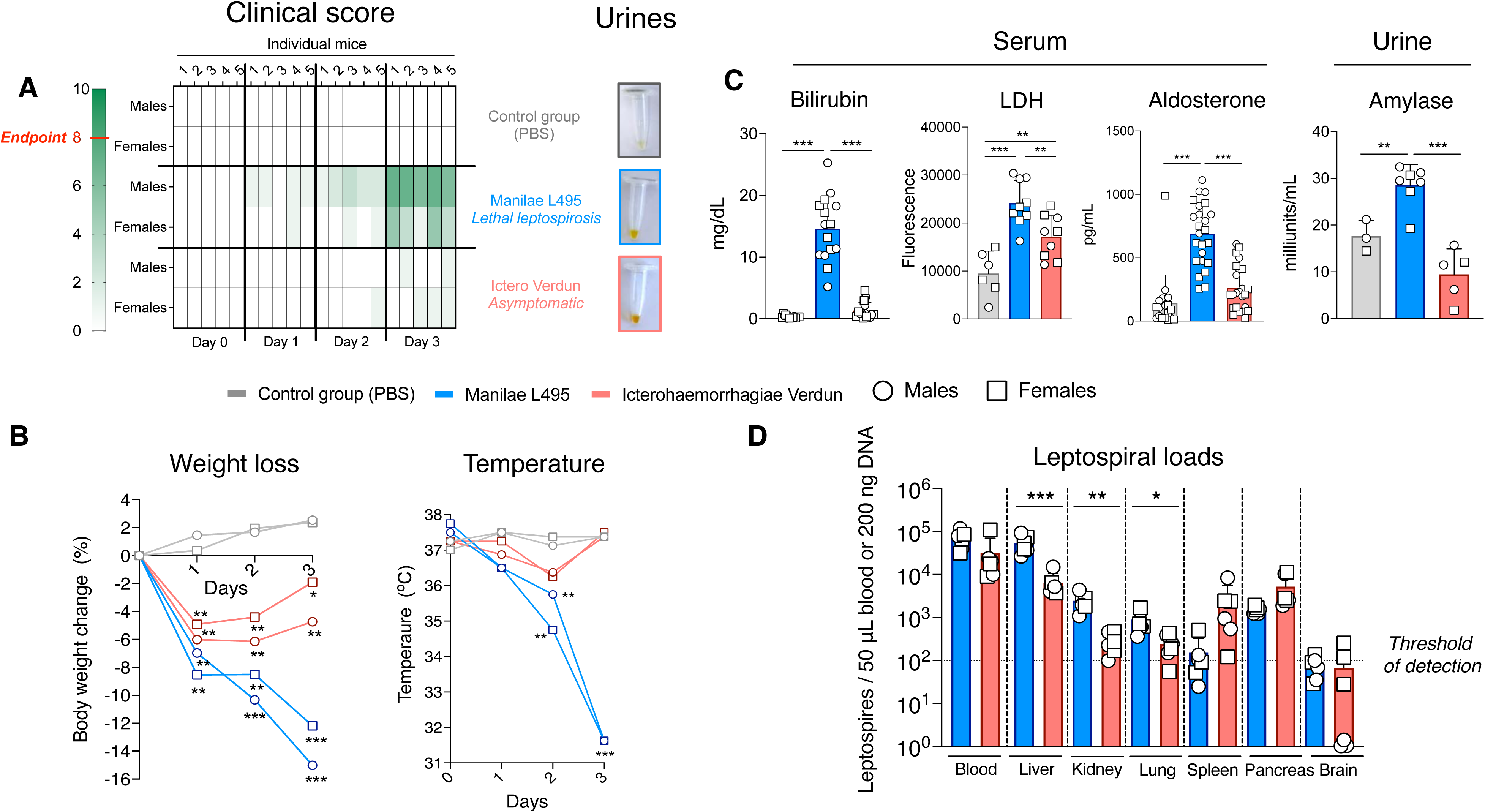
Clinical features of leptospirosis. **(A)** *Left panel*: Table showing the individual clinical scores in both male and female mice infected with *Leptospira interrogans* Manilae L495 and Icterohaemorrhagiae Verdun, or injected with PBS (control group), according to the days post-infection, with color grading corresponding to severity. In dark green, the endpoint 8 corresponds to the humane ethical endpoint. Data represent n=5 mice per group, representative of 3 independent experiments; *Right panel*: Urines of mice collected at day 3 p.i. **(B)** Body weight (n=10 per group, representative of 10 independent experiments) and temperature changes (n=4 per group, representative of 3 independent experiments) in both males (open dots) and females (open squares) during leptospirosis. **(C)** Biochemical markers in serum and urine of mice infected or not with *Leptospira*. Serum levels of bilirubin (n=15), lactate dehydrogenase (LDH) (n=6 for non-infected and n=9 for infected mice), aldosterone (n=18 for non-infected and n=24 for infected mice) and urine dosage of amylase (n=3 for non-infected, n=7 for L495 and n=5 for Verdun-infected mice). **(D)** Leptospiral loads in blood and organs as quantified by qPCR. The limit of detection is set to 10^2^ leptospires per 50 μL of blood or 200 ng DNA. No leptospires were detected in the control group (PBS)(not shown) (n=6 per group, representative of 3 independent experiments).

In conclusion, infection of C57BL/6J mice with 10^8^ *L. i* Manilae strain L495 causes acute severe leptospirosis that results in death of mice with a clinical phenotype resembling sepsis.

### *Leptospira* cause systemic infection

To determine whether leptospires were present in organs, infected mice were sacrificed at the peak of infection (day 3 p.i.) and blood, liver, kidney, lung, spleen, pancreas, and brain were collected. Leptospiral loads in organs were quantified by qPCR of leptospiral DNA. Leptospiral strains were present in each organ. A higher number of leptospires was detected in circulating blood, followed by liver, kidney, pancreas, spleen and lung (Fig. 1D). These results indicate that although the two *Leptospira* serovars induce different clinical symptoms in mice, both are disseminated to all major organs and cause systemic infection. No difference in bacterial load was observed between the two strains in blood, spleen, pancreas and brain, but interestingly, 5 to 10 times more Manilae L495 compared to Icterohaemorrhagiae Verdun were found in lung, liver and kidney, known to be the major target organs of *Leptospira* and often involved in multiorgan failure (21)(Fig. 1D). To confirm these results, the tissues were immunohistochemically stained with an antibody against LipL21, a major lipoprotein of the leptospiral membrane (4). Leptospiral antigens were detected in all organs examined, with discrete *Leptospira* filaments present in the parenchyma of the tissues. Spiral bacteria were observed in liver, renal interstitium, peri-alveolar space and splenic red pulp (Sup Fig. 1A). No sex difference was observed in *Leptospira* quantification by qPCR or in tissue staining for histology.

### *Leptospira*-infected mice show no major signs of liver, kidney, or lung failure

To decipher how mice infected with the Manilae L495 strain die of acute leptospirosis, we studied their liver, kidney, and lungs at day 3 p.i. in comparison with the organs of Verdun-infected mice.

We first quantified two hepatic biomarkers, alanine and aspartate aminotransferases (ALAT and ASAT, respectively), and LDH at day 3 p.i. Unexpectedly, we found no statistically significant difference when comparing the infected groups with the non-infected groups (Fig. 2A). In addition to the quantitative data, histological analysis with hematoxylin and eosin (H&E) further confirmed these results, showing only mild inflammation in the liver tissues of infected mice (Fig. 2B). Compared to uninfected control mice, immunohistochemical staining with anti-F4/80 antibody showed extensive macrophage infiltration in both infected groups, and labeling with anti-Ly6B2 antibody revealed a discrete infiltration of neutrophils. However, a morphologic distinction of neutrophils was observed, with a round, marginal shape in the case of mice infected with Manilae L495, whereas they were diffuse and irregular in the case of Icterohaemorrhagiae Verdun (Fig. 2B).

**Figure 2.**
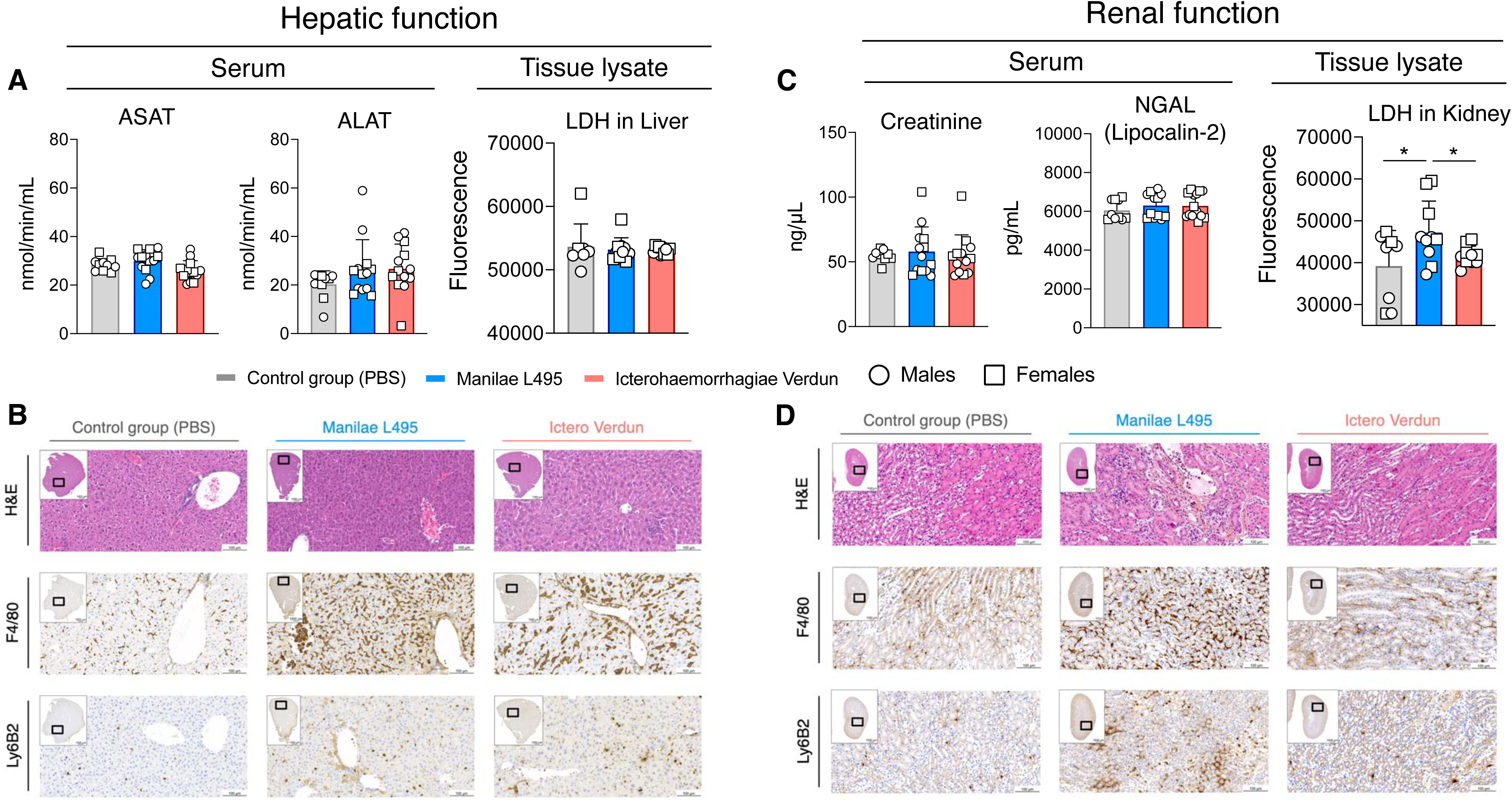
Characteristics of hepatic and renal function in leptospirosis. **(A)** Biochemical markers. Aspartate aminotransferase and alanine aminotransferase (ASAT and ALAT respectively, n=10 for non-infected and n=13 for infected mice for each marker) dosed in serum and lactate dehydrogenase (LDH) n=8 for non-infected and n=10 for infected mice) dosed in the tissue lysate of mice infected or not with *Leptospira*. **(B)** Histology staining of liver tissue of mice infected or not with *Leptospira*. Hematoxylin-Eosin (H&E) staining to detect inflammation, F4/80 to detect macrophages, Ly6B2 to detect neutrophils. The pictures are representative of 3 independent experiments. **(C)** Biochemical markers. Creatinine (n=10 for non-infected and n=13 for infected mice) and Neutrophil gelatinase-associated lipocalin (NGAL or Lipocalin-2, ELISA assay, n=10 for non-infected and n=12 for the *L. i* Manilae L495 and n=15 for the *L. i* Icterohaemorrhagiae Verdun infected mice) dosed in serum and Lactate dehydrogenase (LDH) (n=8 for non-infected and n=10 for infected mice) in the tissue lysate of mice infected or not with *Leptospira*. **(D)** Histology staining of liver tissue of mice infected or not with *Leptospira*. Hematoxylin-Eosin (H&E) staining to detect inflammation, F4/80 to detect macrophages, Ly6B2 to detect neutrophils. The pictures are representative of 3 independent experiments. Open dots represent males and open squares represent females.

Renal function is frequently affected in leptospirosis, as it is associated with severe acute kidney injury in human disease and is also implicated in the chronicity of the infection, as leptospires colonize the proximal renal tubules (23). Therefore, we measured the levels of creatinine and neutrophil gelatinase-associated lipocalin (NGAL), two biomarkers of renal dysfunction, in the blood serum of mice on day 3 p.i. (24). Interestingly, we observed no difference between non-infected and *Leptospira*-infected mice (Fig. 2C). H&E staining of tissues revealed mild hemorrhage only in the kidneys of Manilae L495 infected mice. Anti-Ly6B2 and anti-F4/80 immunostaining showed the recruitment of neutrophils and macrophages upon infection with Manilae L495, but their complete absence in the case of Icterohaemorrhagiae Verdun (Fig. 2D). Consistent with the higher cell infiltration and leptospiral load, the L495 group showed slightly higher LDH release compared to the control or Verdun infected groups (Fig. 2C).

Next, we examined the lungs by immunohistochemistry, since pulmonary hemorrhage is one of the most common manifestations of severe human leptospirosis caused by *L. i*. The results of H&E staining showed signs of mild inflammation upon *Leptospira* infection, with the lungs of the Manilae L495 group being slightly more inflamed compared to those of the Verdun group (Fig. 3A). More neutrophil and macrophage infiltrates were observed in the group of Manilae infected mice (Fig. 3A). Accordingly, the mRNA expression of *inos*, which is responsible for the production of nitric oxide, an antimicrobial compound also known to be damaging to tissues, was increased only in the lethally infected mice. However, the lungs of all groups showed no increase in LDH release. These results suggest that pulmonary inflammation is not sufficient to cause respiratory distress in L495-infected mice.

**Figure 3.**
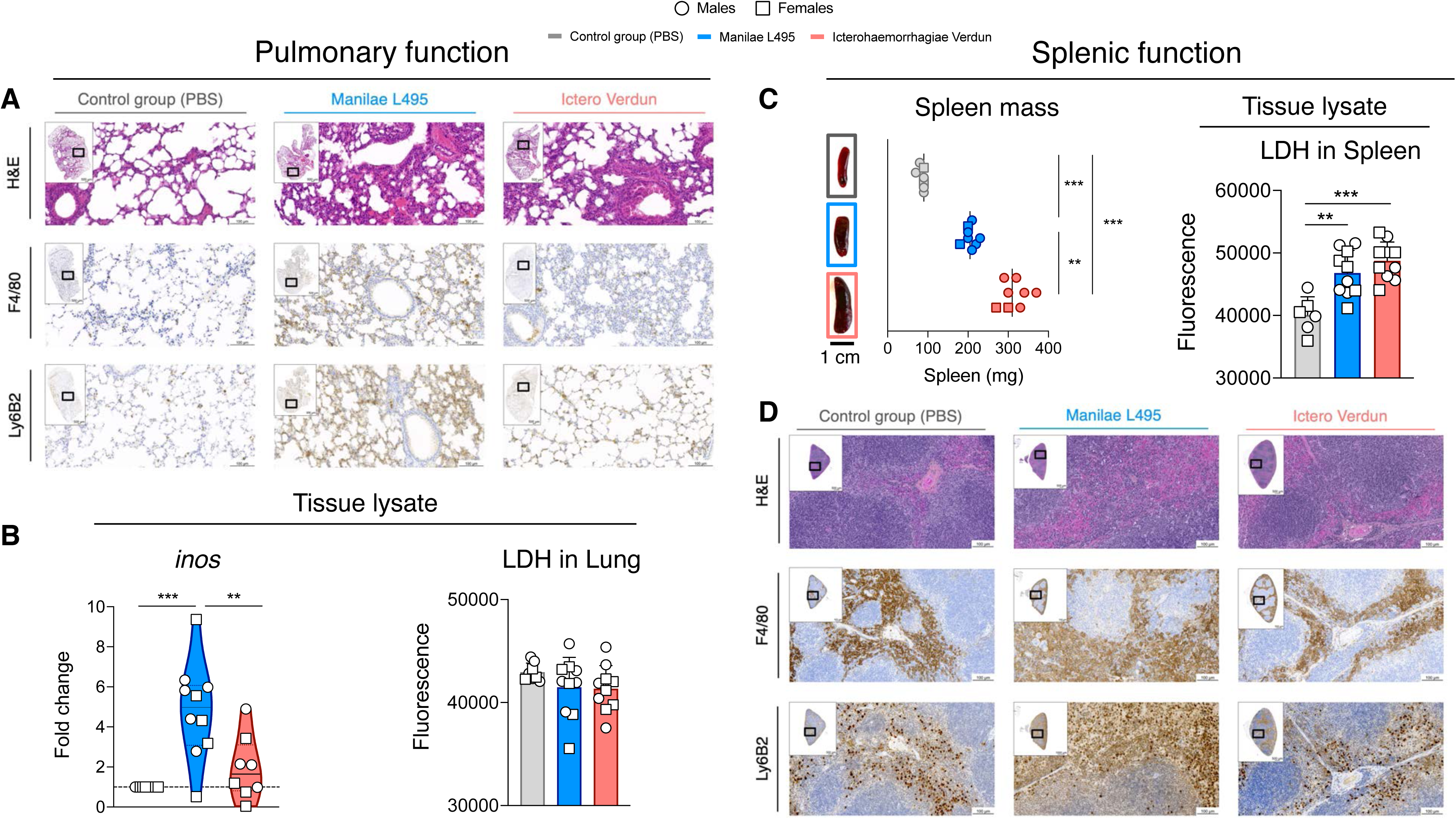
Characteristics of lung and spleen function in leptospirosis. **(A)** Histology staining of lung tissue of mice infected or not with *Leptospira*. Hematoxylin-Eosin (H&E) staining to detect inflammation, F4/80 to detect macrophages, Ly6B2 to detect neutrophils. The pictures are representative of 3 independent experiments. **(B)** Markers in lung lysate. *Inos mRNA* expression fold changes (n=10 for non-infected and L495 infected mice and n=8 for Verdun infected mice) and Lactate dehydrogenase (LDH) (n=8 for non-infected and n=10 for infected mice) in the tissue lysate of mice infected or not with *Leptospira*. **(C)** Characteristics of the splenic function. The mass of the spleen was calculated upon weighting of the whole organ right after dissecting the mice (n=7 for non-infected and n=8 for infected mice). The scale bar below the photos corresponds to 1 cm. Lactate dehydrogenase (LDH) (n=6 for non-infected and n=10 for *Leptospira* infected mice) in the tissue lysate of mice infected or not with *Leptospira*. **(D)** Histology staining of spleen tissue of mice infected or not with *Leptospira*. Hematoxylin-Eosin (H&E) staining to detect inflammation, F4/80 to detect macrophages, Ly6B2 to detect neutrophils. The pictures are representative of 3 independent experiments. Open dots represent males and open squares represent females.

Although leptospires were disseminated in each of the aforementioned tissues, there was no evidence of significant inflammation to justify the death of the mice. Taken together, our data show that L495 lethally infected mice do not show major signs of hepatic, renal and pulmonary insufficiency despite the systemic dissemination of leptospires.

### Leptospiral infection provokes splenomegaly

We next examined the spleens of infected mice, as we observed obvious splenomegaly in the Verdun group (Fig. 3C). Comparison of tissue weight of infected spleens showed increased mass in the Verdun group and to a lesser extent in the Manilae group, both heavier and darker than the uninfected control group. Correspondingly, LDH release was increased in both infected groups and was found to be higher in the Verdun group (Fig. 3C). Although H&E staining of whole spleens did not reveal any obvious signs of lesions, immunostaining revealed a large infiltration of macrophages in both groups, whereas neutrophils were heavily recruited only in the Manilae group (Fig. 3D).

### Pancreatitis is associated with acute leptospirosis

These observations, combined with the striking clinical manifestations and hypothermia, suggested systemic dysfunction. Since no major liver damage was found to explain the bilirubinemia, nor renal insufficiency to explain the urinary amylase levels, we hypothesized that the pancreas might be damaged, as this is a feature occasionally described as a complication of leptospirosis, associated with intensive care unit (ICU) admission and increased mortality rates (25). Since high serum bilirubin and hyperamylasemia have occasionally been reported in patients with leptospirosis who did not develop acute kidney injury (AKI) but pancreatitis (25), we also measured serum amylase (Fig. 4A). The serum results confirmed the urine results, with elevated levels only in the Manilae group. Finally, since the enzyme lipase is known to have a higher specificity for the diagnosis of pancreatitis, we quantified it in the serum of *Leptospira*-infected mice and found that its levels were slightly elevated only in the Manilae L495 group. Therefore, the pancreases of the infected mice were harvested on day 3 p.i. and stained for immunohistology analysis. Notably, H&E staining revealed edema in the epithelium of infected mice (Fig. 4B). A greater infiltration of macrophages and neutrophils, expected features of pancreatitis, was observed in the tissues of Manilae L495 mice compared to those infected with Icterohaemorrhagiae Verdun (Fig. 4B). In addition, a few apoptotic cells were labeled with caspase 3 antibody in the Manilae L495-infected pancreas (Fig. 4B). Taken together, our data showing slightly elevated serum amylase and lipase, together with the histopathologic features, clearly demonstrate pancreatitis as a feature of acute lethal leptospirosis.

**Figure 4.**
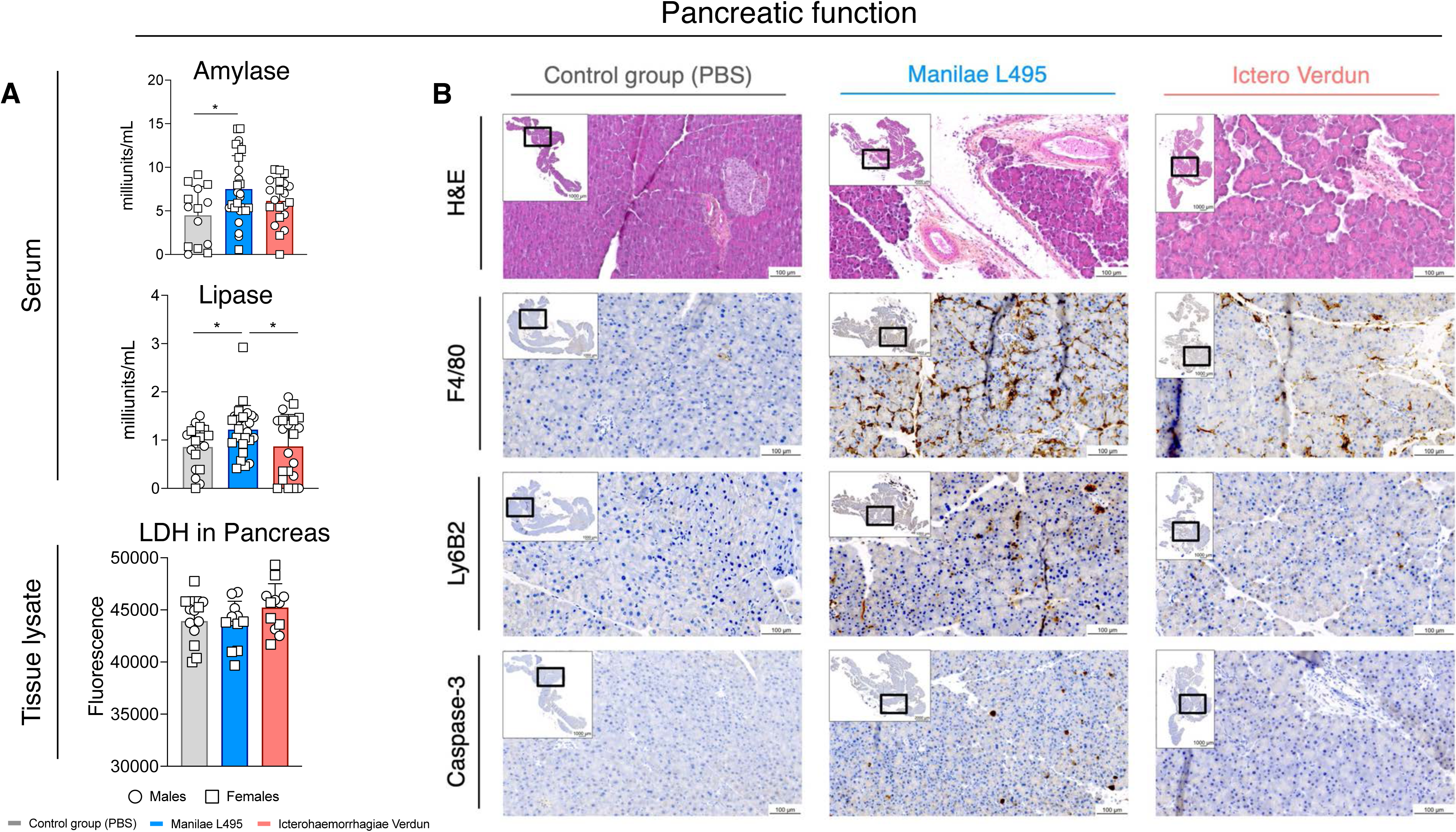
Characteristics of the pancreatic function in leptospirosis. **(A)** Biochemical markers. in serum and liver lysate. Amylase and lipase (n=14 for non-infected, n=27 for L495-infected mice and n=21 for Verdun-infected mice for each marker) in serum and lactate dehydrogenase (LDH) (n=12 for each group) in the tissue lysate of mice infected or not with *Leptospira*. **(B)** Histology staining of pancreas of mice infected or not with *Leptospira*. Hematoxylin-Eosin (H&E) staining to detect inflammation, F4/80 to detect macrophages, Ly6B2 to detect neutrophils and Caspase-3 to detect apoptosis. The pictures are representative of 3 independent experiments. Open dots represent males and open squares represent females.

### No invasive brain infection

Finally, because the death was sudden and because encephalitis with the presence of *Leptospira* in the CSF has been described in human leptospirosis (4), we measured the bacterial loads in the brains (Fig. 1D). Brains from both L495- and Verdun-infected mice showed equal and low numbers of leptospires, barely above the limit of detection. Therefore, we also performed histochemistry in the brain (Fig. 1A, 1B). However, we did not detect any lesions by H&E or LipL21 labeling of leptospires. Since both Verdun- and L495-infected mice showed the same pattern, this suggests that the mice did not die from invasive cerebral infection.

### *Leptospira* do not trigger the burst of a cytokine storm

In our lethal model of acute leptospirosis, mice die from infection with clinical manifestations that mimic septic shock. The production of cytokines, especially TNF and IL-1β, is critical for the initiation of the cytokine storm, generalized inflammation, multi-organ dysfunction syndrome (MODS), and ultimately septic shock and death (26). Therefore, we investigated the pro-inflammatory (TNF, IL-1β, IL-6, IFN-γ) and anti-inflammatory (IL-10) cytokines as well as a chemokine (CCL5/RANTES) in organ homogenates and blood serum of female and male mice on day 3 p.i. by ELISA (Fig. 5). First, the levels of TNF and IL-1β were dramatically low in the spleen, lung and kidney of all infected animals. Strikingly, we observed even less IL-1β in the spleen of both infected groups and in the kidney of the Manilae group. A small increase in TNF and IL-1β was observed only in the liver of Manilae infected mice. IL-6 is an important marker of the severity of sepsis, particularly as an early indicator of the inflammatory state (26). Overall, we did not observe any IL-6 modulation, except for a minimal increase in the kidney of Verdun-infected male mice. IFN-γ is a crucial effector of innate and adaptive immunity, as it primes macrophages and enhances their phagocytic activity. Surprisingly, regardless of the infecting strain, this cytokine was expressed at low levels in all organs, with no statistical difference from control mice. IL-10 is a potent immunosuppressive cytokine whose plasma concentration is increased in sepsis and is associated with adverse outcomes (9). Here, we measured a small increase in IL-10 levels in the livers and lungs of Manilae L495-infected male mice and in the kidneys of Icterohaemorrhagiae Verdun male mice. On the other hand, IL-10 levels were lower in the kidneys (female and male) of the Manilae L495 group than in the uninfected group. Chemokines are important for the recruitment and migration of leukocytes to infection sites. RANTES levels were significantly increased in the spleen of all infected mice, in the kidneys of Manilae L495-infected mice and in Verdun-infected males. Decreased levels were observed in the lungs of males from the Manilae L495 group. In addition, although *Leptospira* induced a systemic infection and leptospiral loads in the circulation were very high on day 3 p.i., we did not find any pro-inflammatory cytokines in the serum of infected mice. However, we measured more IL-10 and RANTES in the serum of all infected mice (Fig. 5). As these results were unexpected, we repeated our experiments with RT-qPCR on RNAs isolated from mouse organs (Fig. 2A), which further confirmed at the transcriptional level the global lack of upregulation of proinflammatory cytokines in organs of infected mice compared to non-infected mice. The only discrepancy observed was *il10* mRNA levels, which were higher in the kidneys of the Manilae group, although the corresponding protein levels were lower. Notably, as in the other organs, we did not measure elevated levels of the cytokines TNF, IL-1β, IL-6, IFN-γ, IL-10 and RANTES in the pancreatic homogenates (Sup Fig. 2B).

**Figure 5.**
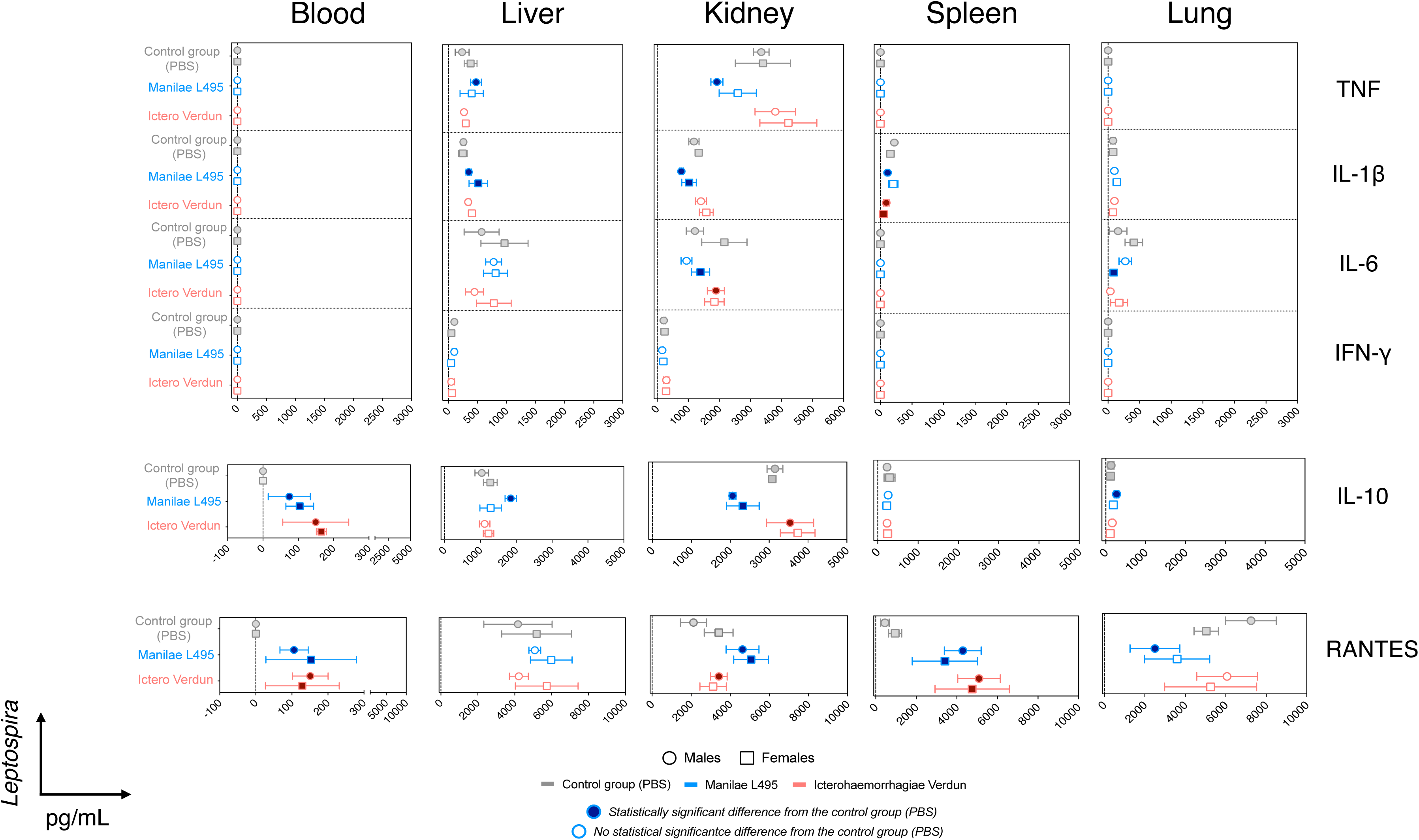
Quantification of cytokines in blood and organs at the peak of the leptospiral infection. Pro-inflammatory TNF, IL-1β, IL-6 and IFN-γ, anti-inflammatory IL-10 and chemokine CCL5/RANTES were measured by ELISA in blood and organs (liver, kidney, spleen, lung) collected at day 3 post-*Leptospira* infection. In the forest plots, each point represents the mean of n=3 non-infected and n=5 *Leptospira* infected mice. Dark bold colors (blue for L495, red for Verdun) represent the statistically significant differences of the infected groups compared to the non-infected one (in grey). Open dots represent males and open squares represent females.

Collectively, our data underscore that despite their sepsis-like phenotype, *Leptospira*-infected mice did not develop hypercytokinemia, but instead displayed increased secretion of the anti-inflammatory cytokine IL-10 and the chemokine RANTES. Thus, lethally infected mice with Manilae L495 did not die from hyperinflammation caused by excessive expression of pro-inflammatory cytokines.

### Septic shock is not induced by *Leptospira*

To completely exclude the hypothesis of a possible septic shock underlying the pathophysiology of lethal leptospirosis, we decided to compare the inflammation induced by injection of inactivated *Leptospira* into mice with that induced by *Escherichia coli* K12, a model bacterium with inflammatory LPS that can be used to experimentally induce septic shock. Because of the difference in generation time between leptospires (18 h) and *E. coli* (20 min), *L. i* Manilae L495, Icterohaemorrhagiae Verdun and *E. c* K12 were heat-inactivated. This stopped their growth and further exposed their MAMPs, mainly LPS. We injected both male and female mice with 10^8^ of the aforementioned heat-killed (HK) bacteria and scored their symptoms as before (Sup Fig. 3A). As expected, mice injected with HK *E. c* K12 developed the clinical signs of septic shock at 4 h p.i. with lethargy, prostration and small eyes, similar to the clinical signs observed on day 3 p.i. in the Manilae infected group (Fig. 1A). On the other hand, HK *L. i* injected mice did not show any signs of suffering or distress during the same period (Sup Fig. 3A), nor did they develop any symptoms later, as previously shown (18). Since the HK *E. c* mice reached the ethical endpoints, we euthanized all mice at 4 h p.i. To investigate inflammation, blood was collected 4 h post-injection and pro-/anti-inflammatory cytokines (TNF, IL-1β, IL-6, IFN-γ, IL-10) along with the chemokine RANTES were quantified in serum by ELISA (Sup Fig. 3B). Interestingly, both HK *L. i* serovars injected to the mice led to the production of very small cytokine levels in blood serum. Notably, all cytokine and chemokine levels were significantly higher in the case of the HK *E. c* K12 group compared to the non-infected one, in both female and male mice, while only RANTES in the HK *L. i* infected mice showed a significant difference compared to control mice (Sup Fig. 3B). These results unambiguously showed that inactivated leptospires per se do not trigger sepsis shock.

However, a recent study showed that severe leptospirosis in hamster alters intestinal barrier integrity (27). To definitively exclude the involvement of a delayed septic shock due to a possible disruption of the intestinal barrier, we collected the small and large intestines of the infected mice at day 3 p.i. Initial observation revealed the absence of food and feces (Sup Fig. 4A and 4B), only in the intestine of the Manilae L495 infected mice. H&E staining confirmed the very few pieces of food in the gut of this group, suggesting that Manilae infection induces anorexia. Histological staining (H&E, anti-LipL21) of the small and large intestines of *Leptospira*-infected mice at day 3 p.i. did not reveal cellular infiltrates, the presence of leptospires, or lesions of the epithelial barrier (Fig. 4A and 4B). To further investigate the integrity of the intestinal barrier in the *Leptospira*-infected mice and the possible translocation of the intestinal microbiota into the blood circulation, we collected their blood at day 3 p.i. and inoculated it into special long tubes containing Brain Heart Infusion (BHI) medium to allow the growth of both anaerobic and aerobic bacteria (Sup Fig. 4C). Interestingly, we didn’t observe any bacterial growth, concluding that the intestinal mucosal barrier is not damaged by *Leptospira* infection.

Taken together, these results are consistent with the absence of cytokine storm and clearly indicate that *Leptospira* does not induce septic shock in our model of lethal leptospirosis, despite the similar and common clinical symptoms.

### Neutrophils are recruited into blood and organs of the lethally infected mice

Our results showed that mouse death is independent of hyperinflammation or cytokine storm, but immunohistochemistry revealed distinct neutrophil and macrophage profiles upon infection with leptospires. To gain more insight on the cellular changes occurring in the lethally infected mice, we next phenotyped by cytometry the total cell populations in the liver, kidney, lung, spleen, and blood at day 3 p.i. (Fig. 6A and Sup Fig. 5A, 5B, 5C). Ly6G^+^ neutrophils were infiltrated in the liver, kidney, and lung of mice infected with *L. i* Manilae L495 but not *L. i* Icterohaemorrhagiae Verdun (Fig. 6A, left panel), demonstrating that lethal infection induces neutrophils to cross the endothelial barrier and migrate into the tissues. In contrast, increased numbers of neutrophils were observed in the blood and spleen of both Manilae and Verdun groups (Fig. 6A, left panel). Ly6C^low^ monocytes were also found to be significantly increased only in Manilae L495-infected livers and lungs, as well as in the circulation of mice infected with Manilae L495 and Icterohaemorrhagiae Verdun (Fig. 6A, right panel). Strikingly, the profile of other cell types, such as Siglec F^+^ eosinophils, CD11b^+^ and CD103^+^ DCs, was not altered in the infected organs (Sup Fig. 5B), except for CD11b^+^ DCs in the liver of Verdun-infected mice. Consistent with the increase in neutrophils and monocytes, the percentage of B and T lymphocytes was significantly decreased in the organs and blood of mice infected with the two *Leptospira* strains (Sup Fig. 5C). Of note, the experiments were performed in mice of both sexes, with no difference between female and male individuals. Taken together, these results demonstrate that the two *Leptospira* strains induce distinct changes in the nature of the circulating cells and in the recruitment of cell populations into the tissues, most likely depending on the severity of the infection. Interestingly, neutrophils were the only subset of myeloid cells that infiltrated all organs tested in Manilae L495 infected mice.

**Figure 6.**
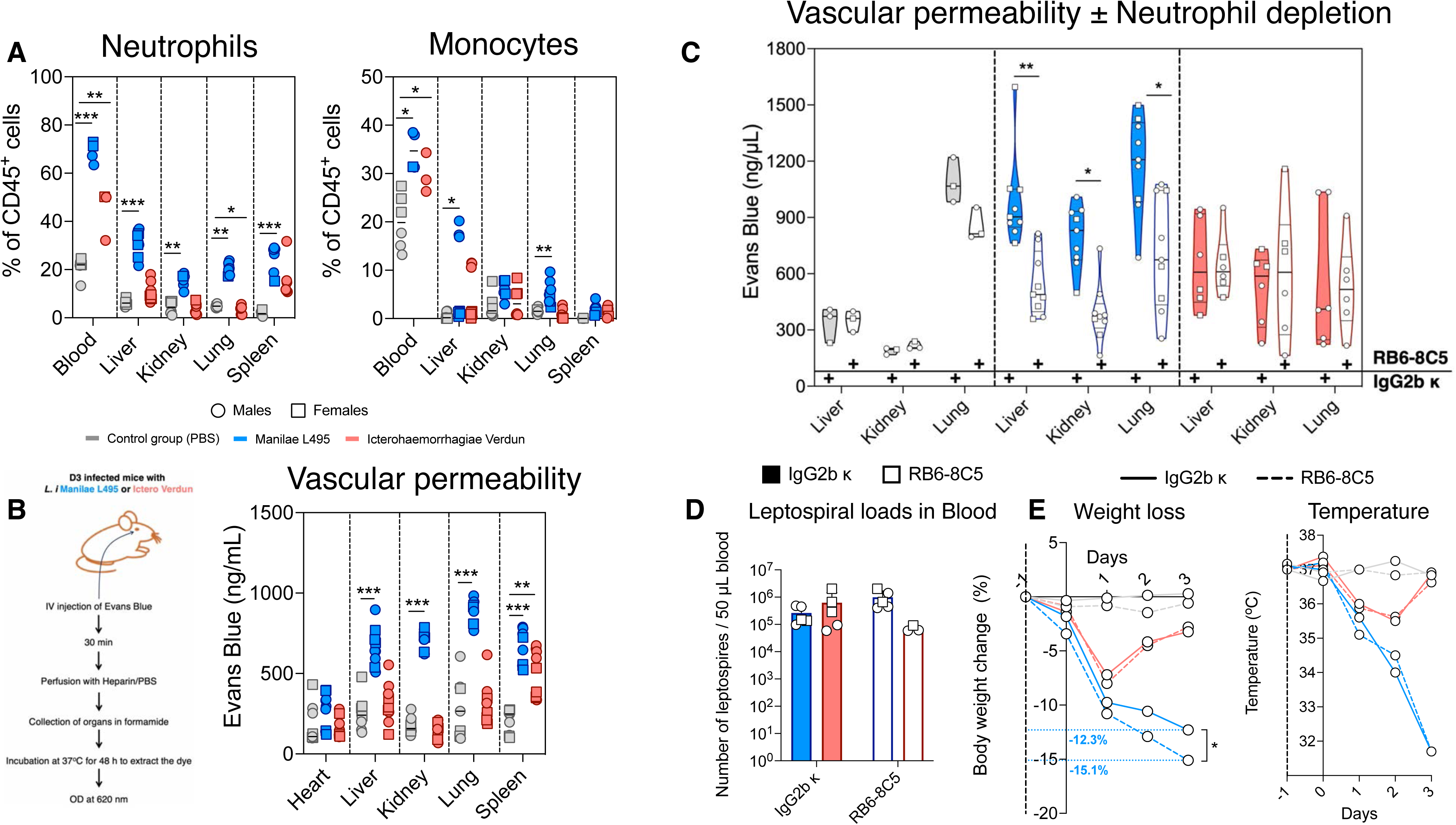
Neutrophils infiltrate during acute lethal leptospirosis and are responsible for vascular permeability. **(A)** Flow cytometry analysis of neutrophils and monocytes in blood and organs (liver, kidney, lung, spleen). Neutrophils (CD11b^+^Ly6G^+^) are recruited in blood and infiltrated in liver, kidney, lung, spleen upon infection with *L. i* Manilae L495 while only in blood and spleen upon *L. i* Icterohaemorrrhagiae Verdun infection. Monocytes (CD11b^high^MHCII^-^Ly6C^high^) are infiltrated in blood upon infection with both *Leptospira* strains and infiltrated in liver upon *L. i* Manilae L495 infection (compilation of n=3 independent experiments). **(B)** *Left panel*: Schematic representation of the protocol used to test the vascular permeability. *Right panel*: Lethal infection with *L. i* Manilae L495 provokes vascular permeability in liver, kidney, lung and spleen as measured by the Evans blue assay. *L. i* Icterohaemorrhagiae Verdun increases the vascular permeability only in spleen. **(C)** Vascular permeability in the presence or absence of neutrophils as determined by the Evans blue assay. Lethal infection with *L. i* Manilae L495 in the absence of neutrophils decreases the vascular permeability. Bold violin plots represent the non-depleted (IgG2b κ) mice while white violin plots represent the neutrophil-depleted (RB6-8C5) ones (compilation of n=3 independent experiments). **(D)** Leptospiral loads in blood in the presence (IgG2b κ) or absence (RB6-8C5) of neutrophils. The loads are not altered upon depletion of neutrophils regardless the *Leptospira* strain used for the infection (representative graph of n=2 experiments). **(E)** Body weight and temperature changes upon *Leptospira* infection in the presence (IgG2b κ) or absence (RB6-8C5) of neutrophils. Neutrophil-depleted mice lost more weight compared to the non-depleted ones. No change was observed in temperature (representative graph of n=3 experiments). Open dots represent males and open squares represent females.

### Lethal leptospirosis induces neutrophil-dependent increased vascular permeability

Since neutrophils are usually the first line of innate immune defense against bacteria and infiltrate all tissues of lethally infected mice, we hypothesize that they may influence the course of acute leptospirosis. Since one of the best studied side effects of neutrophils is endothelial damage (28), we measured vascular permeability using Evans blue (Fig. 6B) (Fig. 5B left schematic). The results show increased vascular permeability at day 3 p.i. in liver, kidney, lung and spleen of the group infected with Manilae L495 strain. In contrast, no vascular permeability was measured in the organs of the Verdun group, except for the spleen (Figure 6B). To verify the involvement of neutrophils in this phenotype, we next proceeded to their depletion with an antibody directed against GR-1 (clone RB6-8C5) one day before infection. This treatment resulted in an effective depletion of circulating neutrophils compared to mice receiving the isotype control antibody, as confirmed by flow cytometry analysis (Sup Fig. 6). Correspondingly, the proportion of monocytes among CD45^+^ cells increased in both groups after neutrophil depletion (Sup Fig. 6). Strikingly, vascular leakage due to Manilae infection was drastically reduced in the absence of neutrophils (Fig. 6C, middle panel), suggesting their potential deleterious role on the endothelium during acute leptospirosis. In contrast, neutrophil depletion in the Verdun group did not alter baseline dye levels (Fig. 6C, right panel). Taken together, these experiments suggest that polymorphonuclear neutrophils have a detrimental effect on the host by causing damage to the endothelial barrier. Interestingly, the leptospiral load measured by qPCR in depleted and non-depleted mice was not significantly different (Fig. 6D), suggesting that neutrophils were not effective in clearing Manilae L495. However, their depletion in Manilae-infected mice did not alter the temperature (Fig. 6E), but instead exacerbated the disease, as observed by the slight increase in weight loss compared to non-depleted mice (Fig. 6E). Since neutrophil depletion did not attenuate the infection, we speculated that the death of Manilae-infected mice was due to reasons other than vascular leakage, leading us to speculate on a vital organ that might be involved, such as the heart.

### Myocarditis in lethal leptospirosis

Heart problems in patients with acute leptospirosis are mentioned in some case reports (29, 30), which led us to examine the hearts of infected mice at day 3 p.i. First, we measured the heart weight, as it is a reliable marker of heart disease in mice (31). Interestingly, unlike the hearts of Verdun-infected mice, those of the Manilae group were heavier than those of uninfected mice (Fig. 7A). We also measured two markers in the serum of *Leptospira*-infected mice, both of which are typical clinical signs of myocarditis: cardiac troponin I (cTnI) and creatine kinase-myocardial band (CK-MB). Indeed, cTnI and CK-MB levels were elevated in the serum of the Manilae L495-infected group but not in that of the Icterohaemorrhagiae Verdun-infected group (Fig. 7B), suggesting cardiac dysfunction in lethally infected mice. However, no LDH elevation (Fig. 7C) or caspase-3 histologic labeling (Sup. Fig. 2C) was detected in the hearts of *Leptospira*-infected mice, showing that cell death is not involved in heart dysfunction. Cardiac tissue was then examined by immunohistology. As expected, leptospires labeled with anti-LipL21 were abundant in the blood vessels, in accordance with the bacterial load found in the circulation (Fig. 1D). However, very few of them were located in the myocardium near the vessels (Sup Fig. 1). H&E staining revealed cellular infiltrates in the myocardium of Manilae-infected mice, which were characterized as macrophages by anti-F4/80 labeling and as T cells by anti-CD3 antibody. Interestingly, both macrophage and T-cell infiltrates are histological features of myocarditis (32). In the Verdun group, macrophages were also observed in the myocardium, but not T cells, and neutrophils were only observed in the circulation, but not in the myocardium (Fig. 7D). Consistent with the phenotyping of cell populations in the blood (Fig. 5A), higher numbers of neutrophils were observed in the vessels of mice infected with Manilae L495 compared to those infected with Icterohaemorrhagiae Verdun. The infiltration of macrophages and T lymphocytes led us to check for a possible ongoing inflammation in hearts. Cytokines were quantified by ELISA in hearts homogenates, but similarly to livers, kidneys, lungs, spleens and blood, no proinflammatory cytokines (TNF, IL-1β, IL-6, IFN-γ) were detected (Sup Fig. 2C). Anti-inflammatory IL-10 levels were measurable but did not show any statistically significant increase compared to the PBS-treated mice. However, an increase of RANTES was observed in the infected groups compared to the non-infected one (Sup Fig. 2C). These data clearly highlight the absence of inflammation in the hearts of the infected mice. We did not observe any sex differences in the development of the cardiac dysfunction. Taken together, our data reveal an unappreciated and critical role of myocarditis associated with lethal leptospirosis, contributing to the death of mice.

**Figure 7.**
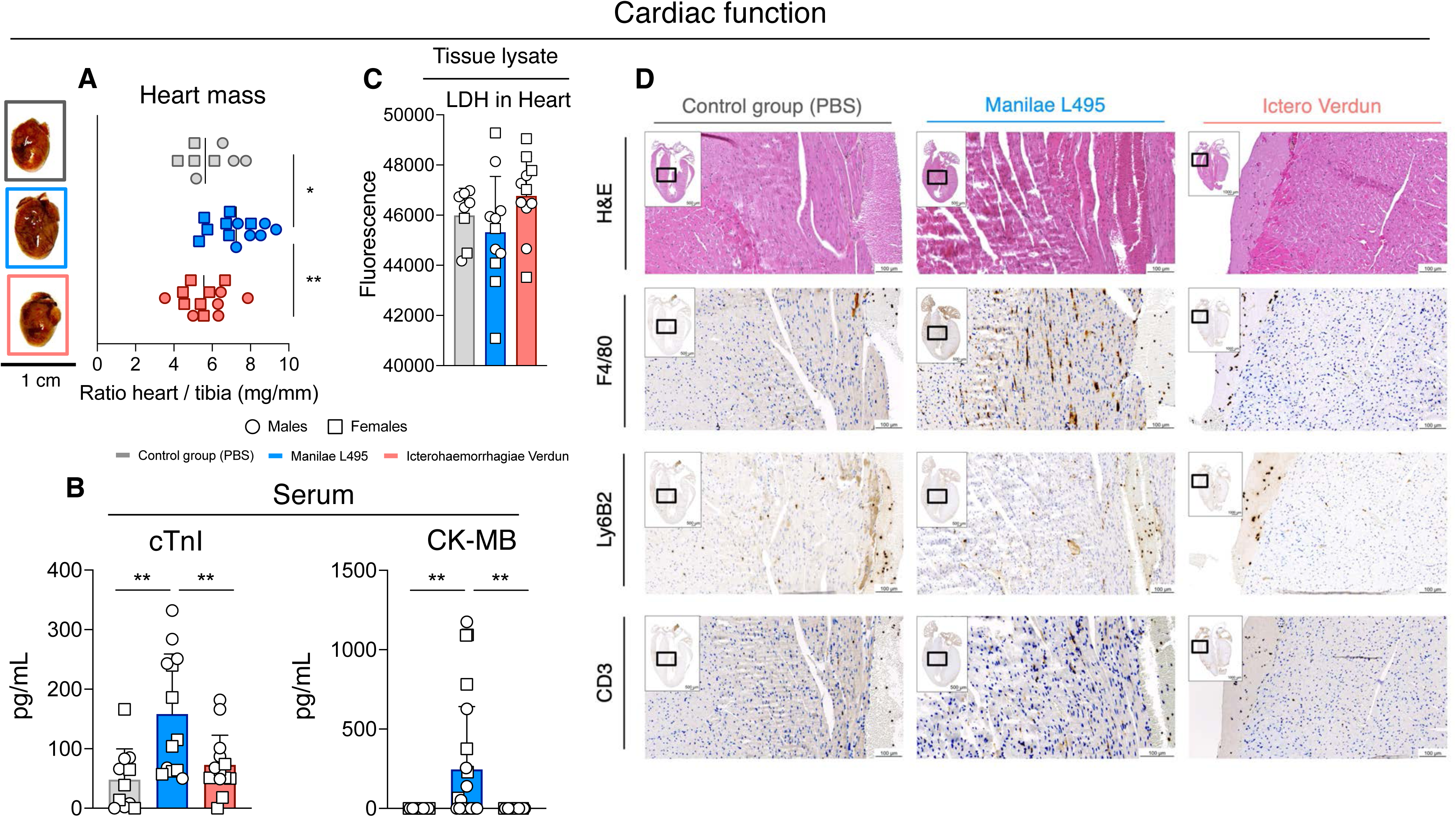
Characteristics of the cardiac function in leptospirosis. **(A)** Weight of the whole hearts right after dissecting the mice, further relativized by the length of the tibia of the corresponding mouse (n=8 for non-infected and n=13 for *Leptospira*-infected mice). The scale bar below the photos corresponds to 1 cm. **(B)** Lactate dehydrogenase (LDH) (n=8 for non-infected and n=11 for *Leptospira* infected mice) in the tissue lysate of mice infected or not with *Leptospira*. **(C)** Biochemical markers. Cardiac troponin I (cTnI) (n=11 for non-infected and n=14 for *Leptospira* infected mice) and Creatinine kinase myosin band (CK-MB) (n=22 for non-infected and Verdun infected mice and n=24 for L495 infected mice) were dosed in serum of mice infected or not with *Leptospira*. **(D)** Histology staining of cardiac tissue of mice infected or not with *Leptospira*. Hematoxylin-Eosin (H&E) staining to detect inflammation, F4/80 to detect macrophages, Ly6B2 to detect neutrophils and CD3 to detect T lymphocytes. The pictures are representative of 3 independent experiments. Open dots represent males and open squares represent females.

In conclusion, by studying a novel model of acute leptospirosis in mice, we highlighted unexpected and unanticipated pathophysiological features that explain the death of mice during severe leptospirosis. In fact, we showed that the mice did not die of inflammation but showed splenomegaly and signs of pancreatitis. We showed that the main cause of death was myocarditis, exacerbated by neutrophil-associated vascular damage. These features are also found in patients with leptospirosis, making this study a paradigm for a better understanding of leptospirosis, a neglected and re-emerging zoonosis.

## Discussion

In this work, we characterized a mouse model of acute severe leptospirosis caused by *L. interrogans* serovar Manilae strain L495 (17). A side-by-side comparison of lethal infection with *L. i* serovar Icterohaemorragiae strain Verdun, which did not induce severe disease, helped us to decipher the differential features leading to mouse death (Table 5). Thus, severe leptospirosis is associated with hypothermia, weight loss, elevated levels of bilirubin, aldosterone, amylase, lipase, cardiac troponin I and creatine kinase, markers of pancreatitis and cardiac disease. We also investigated the pathophysiology in the organs using immunohistochemistry and cellular phenotyping, which revealed vascular permeability in the lungs, kidneys and liver, and myocarditis as evidenced by macrophage and T cell infiltrates in the myocardium.

The present results, supported by numerous replications and complementary approaches, are provocative in several respects: i) mice are generally considered “resistant hosts” to leptospirosis; ii) we found no evidence of major lesions in the usual target organs, such as liver, kidney, and lung, although we observed features of acute severe leptospirosis, such as systemic infection with neutrophilia associated with morbidity and jaundice; iii) we did not measure pro-inflammatory cytokines, thus excluding septic shock due to a cytokine storm, despite neutrophil and macrophage infiltration in most organs and a marked anti-inflammatory IL-10 and chemokine response; iv) unexpectedly, we found that the main cause of death in the mice was myocarditis, associated with vascular leakage and pancreatitis, all known but often overlooked complications of human leptospirosis (16, 25, 30).

### First model of lethal leptospirosis using immunocompetent mice

Common models of human leptospirosis are 3-to 4-week-old guinea pigs or Golden Syrian hamsters, which are not sexually mature and lack a competent immune system (33, 34). Previous mouse models of lethal leptospirosis have relied on immunodeficient mouse strains: such as juvenile C3H/HeJ, which naturally carry a nonfunctional *tlr4* mutation, adult C57BL/6J *tlr4^-/-^* transgenic mice, or µMT mice deficient in B cells (33). Often, the bacterial model used was *L. i* serovar Copenhageni, strain Fiocruz L1-130, which does not induce death in wild-type C57BL/6J mice, even at a dose of 5×10^8^ bacteria/mouse. These studies highlighted the critical role of B lymphocytes and TLR4 in mouse defense against *Leptospira* (35). However, all these young or immunodeficient animals could not be considered as good models to study the physiopathology associated with death. Here, using the more virulent *L. i* Manilae L495 strain, which may be better adapted to the mouse than the Icterohaemorrhagiae Verdun strain commonly found in rats (36), we showed that wild-type adult C57BL/6J mice were severely affected by leptospirosis. Several recent studies have indicated that the severity of leptospirosis depends on the serovar and correlates with the bacteriemia (24, 37). Here, we only found a difference in bacterial loads between the *Leptospira* strains in organs that were not at the origin of death, suggesting that other factors are involved in lethality. The more severe cases of human leptospirosis usually affect men (16) which led us to perform our study with both sexes of mice in parallel, which to our knowledge has never been done before in our field. Interestingly, as previously shown in the adult hamster model (38), we found that male mice were slightly more susceptible to lethal infection than females. We anticipate that our mouse model of lethal leptospirosis will be relevant for further investigation of this pronounced sex bias in human leptospirosis, mostly affecting men (4). In addition, this model could be used to better understand the virulence factors involved in severity, for example by comparing the transcriptomic profiles in mouse blood of our two serovars of *Leptospira interrogans*.

### No inflammation

Mice infected with leptospires present an overall systemic increase in the chemokine RANTES which can attract neutrophils (39, 40), but this was not associated with over-production of pro-inflammatory cytokines. This was not expected considering that bone marrow derived macrophages (12) or dendritic cells (14) produce both chemokines and pro-inflammatory cytokines upon infection with leptospires. However, these data are consistent with a recent study from our group showing that escape from pyroptosis by leptospires prevents massive IL-1β release and that IL-1β does not play a major role during acute severe leptospirosis in mice (12). The consequence of the lack of IL-1β release would be to limit the production of other cytokines such as IL-6. This was observed not only in the blood but also in the organs, suggesting an active mechanism of Manilae L495 to dampen pro-inflammatory cytokines but not chemokines. Notably, the levels of pro-inflammatory cytokines in L495-infected kidneys were even found to be significantly lower than in controls. Finally, the elevated levels of the anti-inflammatory cytokine IL-10 in the blood suggest its involvement in the absence of inflammation. Consistent with this finding, studies in mice and hamsters have already shown that IL-10 is an important player in leptospirosis, possibly regulating host resistance and leptospiral persistence (41, 42).

The involvement of cytokines in human leptospirosis is more controversial (7). In fact, the chemokine RANTES and the cytokine IL-6 are usually detected in all patients with leptospirosis and elevated levels correlate with the severity of leptospirosis (24, 43, 44), but discrepant data exist in the field on the role of TNF, IL-1β and IL-10. In addition, IL-1β and TNF cytokines which are hallmark of sepsis are not always detected in patients with complications (44). Moreover, most studies have compared mild to severe leptospirosis and usually find more cytokines in patients with severe disease, but the cytokine levels are not compared to a healthy group, nor to a cohort of patients with sepsis due to other bacteria. It is also important to note that all these data were obtained in patients treated with antibiotics, which can provoke the Jarisch-Herxheimer reaction, with increased cytokine production due to the release of bacterial components that amplify inflammation (45). Therefore, the common view that acute severe leptospirosis is associated with “a cytokine storm” (4, 7) should be carefully reconsidered, as it may mislead clinicians about the true cause of severe disease.

### Neutrophils and Vascular Permeability

Neutrophilia is a known feature of (46) leptospirosis in humans and a predictor of severe disease (37, 47–49). It was hypothesized that neutrophils were unable to migrate into inflamed tissues because the expression of characteristic activation markers was not elevated in patients with leptospirosis (50, 51). However, in our mouse model of lethal leptospirosis, we demonstrated neutrophil recruitment in the blood, consistent with the high levels of RANTES and their infiltration of all major organs typically affected by leptospires. Interestingly, immunohistochemistry of splenic necropsies from deceased patients with leptospirosis also showed moderate to intense infiltration of plasma cells and neutrophils (52), suggesting that human neutrophils are indeed found in infected organs. This also suggests that leptospires do not block their rolling and extravasation into tissues.

We performed neutrophil depletion and demonstrated for the first time the critical role of neutrophils in vascular permeability in leptospirosis. The common belief in the field is that *Leptospira*, through their numerous toxins and proteases, directly affect the endothelium, causing hemorrhage and vascular permeability. Indeed, a recent study showed that *Leptospira* can induce morphological changes and reduce cell-cell junction protein signaling in human endothelial cells (53). Notably, we did not observe hemorrhage in our lethal model, although blood was observed in the urine of mice infected with the Icterohaemorrhagiae Verdun strain, which showed no vascular permeability except in the spleen. This suggests that the deleterious mechanisms affecting the vasculature may differ from one *Leptospira* serovar/strain to another and most likely occur through different mechanisms, either intrinsic to the bacteria or related to neutrophils. Interestingly, leptospires have been shown to activate NETs (54), which could further damage the endothelium through the release of reactive oxygen species and proteases such as elastase. Further studies are planned in mice with human dermis xenografts, thus providing the human vasculature to study the role of neutrophils in endothelial damage using infections with different serovars of *Leptospira* (28).

Although neutrophils are traditionally considered to be the predominant cells at the forefront of the innate defense, their depletion did not alter leptospiral loads in blood or the mice body temperature, suggesting their inefficient ability to control *L. i* Manilae L495. Paradoxically, their depletion even aggravated the weight loss, suggesting that neutrophils still provide some little help to mice in fighting leptospirosis, as previously shown for *L. i* Copenhageni Fiocruz infected mice (55). Indeed, we may hypothesize that NETs that were shown to control, at least partially, the bacterial burden in kidneys of *L. i* Copenhageni Fiocruz infected mice (55) could also be efficient to trap *L. i* Manilae L495 and avoid their dissemination. Further studies are required to study the complex roles of neutrophils (56) in our lethal model of leptospirosis.

Neutrophils also remain a puzzling cell population in Lyme disease, with some studies showing their effective role against *Borrelia* and others showing their inability to fight the bacteria. (57, 58). Interestingly, alteration of the vascular permeability has also been observed in the cases of *Borrelia burgdorferi* and *Treponema pallidum* (57, 59). These results clearly highlight the complex involvement of neutrophils that globally seems deleterious in acute spirochetosis. Our data suggest that therapeutic strategies aimed at controlling neutrophilia may be useful in limiting vascular permeability in patients with severe leptospirosis.

### Reasons of death

Despite advances in genetically engineering *Leptospira*, the host immune response to infection and subsequent clinical manifestations remains largely unexplored, as the precise mechanisms leading to death have yet to be elucidated. The ability of leptospires to efficiently evade recognition by most innate immune mechanisms gives them a silent and discrete profile (10), which, at least in our lethal model, is consistent with the absence of systemic hyperinflammation despite their global spread.

We did not find elevated levels of LDH in liver, lung, heart nor pancreas suggesting that leptospires do not induce pyroptosis or necroptosis in these organs, consistent with our previous work on macrophages (12). However, LDH was elevated in kidney of L495 and in the blood and spleen of both L495- and Verdun-infected mice, consistent with a recent article showing that necroptosis occurs at day 3 p.i. in the spleen of TLR4-deficient mice infected with *L. interrogans* serovar Copenhageni (46). However, together with the lack of inflammation, this suggests that inflammatory cell death is not associated with lethality in our model.

In the present study, we found evidence of pancreatitis, with slightly elevated serum amylase and lipase concentrations, mild pancreatic epithelial edema, and infiltrating macrophages and, to a lesser extent, neutrophils, which are known to be important cellular players in pancreatitis (60). Indeed, as mentioned above, NETs are involved as a defense mechanism against *Leptospira*, but they can also be harmful by promoting pancreatic tissue damage in various ways (60). Pancreatitis has long been recognized as a complication in patients with leptospirosis, and several individual case reports have been published over the years (61–64). A recent systematic review (25) summarizes the clinical features of acute pancreatitis in leptospirosis and suggests that this overlooked complication of the disease may not be so rare. When serum amylase and lipase were measured in people with acute leptospirosis, their levels increased more than threefold compared to reference values (49), which is higher than what was measured in our model. Therefore, the absence of LDH release and the low number of apoptotic cells, together with the only slight increase of the markers, indicate the mildness of the pancreatitis in our model of mice infected with *L. i* Manilae L495. These observations suggest that the possibility of mice dying from pancreatitis remains low. Nevertheless, the bilirubinemia measured in the lethally infected mice, possibly due to compression of the bile ducts by pancreatic damage, could be related to the elevated cardiac troponin I levels and consequent cardiac dysfunction. Indeed, it has been proposed that total bilirubin levels are associated with an increase in troponin, which is the best indicator of myocardial injury in COVID-19 (65) and, additionally, troponin was also found elevated in patients with pancreatitis (66). Thus, the implication of pancreas in acute leptospirosis might be rather indirect than direct.

The main feature responsible for mouse death in our model is myocarditis, as lethally infected mice develop its typical symptoms, including an increase in troponin and CK-MB, as well as macrophage and T cell infiltration of the cardiac muscle (67). The involvement of myocardial damage in leptospirosis is extremely neglected, underdiagnosed and overlooked in most patients. Another element supporting myocarditis is the association between global myocardial damage and elevated aldosterone levels, which are also associated with hypertension (68) and disruption of the sodium/potassium equilibrium (69). Of note leptospires are known to inhibit the Na^+^/K^+^ ATPase pump, and other transporters in kidneys, leading to potassium efflux and inflammasome activation (70). In addition, a recent study correlated increased blood aldosterone concentration with increased myoglobin, a marker of cardiac injury (71). Because arrhythmias have been observed in human leptospirosis (30) and because sudden death occurs early in infection, it would be interesting to study this in our lethal model. Unfortunately, monitoring cardiac parameters and echocardiography in mice is complicated and we do not have access to this extensive equipment in our infectious animal facility.

Very few case reports (29, 30, 72) mentioned cardiac manifestations as a complication of leptospirosis, always associated with MODS and high levels of markers associated with myocardial injury, a fact that clearly indicates the prevalence of cardiac injury during the acute phase of the disease. However, this is not so rare, as in Sri Lanka, 15% of people infected after a flood were diagnosed with myocarditis (73). In French West Indies (Guadeloupe) it was recently shown that 41 % of critically ill patient with leptospirosis had myocarditis which was significatively associated with death at the ICU (74).

However, the general interest of our model may not be limited to the case of leptospirosis. Myocarditis is also considered a rare consequence of Lyme disease in humans (75). The study of myocarditis using experimental murine models can be complicated and the experimental diseases are typically induced by cardiotropic agents or by activating autoimmune responses (32). However, the etiologies of myocarditis in humans can be either non-infectious or infectious, with viruses such as HIV, EBV (76) or COVID-19 (77) being some of the most dominant agents. In addition, the protozoan parasite *Trypanosoma cruzi* can also induce myocarditis in mice (78). Thus, our lethal model of leptospirosis not only reflects the manifestations of human disease in the acute phase of leptospirosis but could also be used as a paradigm for further study of microbe-induced acute myocarditis.

In summary, although we must take into account the host species specificity of leptospirosis (10) and the broad clinical manifestations of the disease (4), our lethal mouse model accurately mimics several features of human leptospirosis, in particular neutrophilia, vascular damage, pancreatitis, and myocarditis. Importantly, this work rules out the cytokine storm as a cause of death.

## Acknowledgments

We thank Tiphaine Camarasa and Mélanie Hamon (Institut Pasteur) for help in lung phenotyping. We thank Delphine Bonhomme for help in the sepsis model. We thank Jean-Marc Cavaillon for critical reading of the manuscript. We thank Pr. Jean-Pierre Hulot (Hôpital Européen Georges-Pompidou, Paris, France) and Timothy Wai (Institut Pasteur) for discussions about myocarditis and heart disease. We thank Dr. François Baur (Poindimié) and Dr. Patrick Lefevre (Koumac) from the North Hospitals of New Caledonia for fruitful discussions about human leptospirosis. We thank Dr. Amaro Duarte (Faculty of Medicine, University of São Paulo, Brazil) for discussions about histochemistry of organs of deceased patients with leptospirosis. We thank members of the Histopathology Core Facility at Institut Pasteur, for helping with the sample preparation and staining. We acknowledge the Flow Cytometry platform at Institut Pasteur for support in conducting this study. CW used “DeepL write” for English edition. After using this tool, CW, SP and all authors reviewed and edited the content and take full responsibility for the content of the publication.

## Funding

The Boneca laboratory was supported by the following programmes: Investissement d’Avenir program, Laboratoire d’Excellence “Integrative Biology of Emerging Infectious Diseases” (ANR-10-LABX-62-IBEID) and by R&D grants from Danone and MEIJI. CW received an ICRAD/ANR grant (S-CR23012-ANR 22 ICRD 0004 01).

SP received a scholarship by Université Paris Cité (formerly Université Paris V – Descartes) through Doctoral School BioSPC (ED562, BioSPC). SP has additionally received a scholarship “Fin de Thèse de Science” number FDT202404018322 granted by “Fondation pour la Recherche Médicale (FRM)”.

## Authors contributions

Conception and supervision of the project: CW Investigation, validation and figures: SP Methodology: SP, MT, FVP Data analysis: SP, DH and CW Funding acquisition: CW and IGB Writing: SP and CW All authors contributed to the review and editing of the manuscript and approved it for submission.

---

**Supplementary Figure 1.**
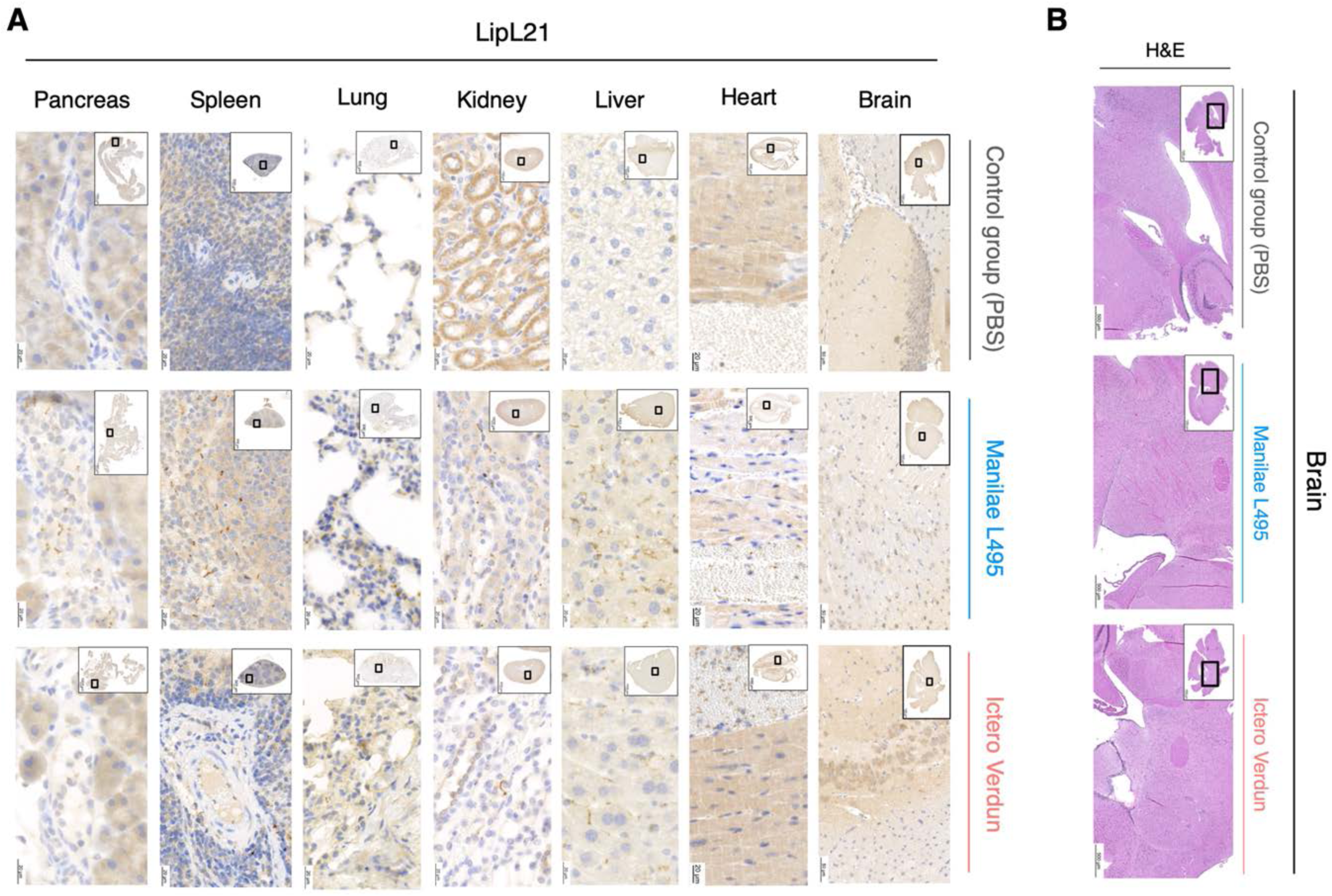
Detection of *Leptospira* with immunohistochemical staining of anti-LipL21 and absence of brain inflammation. **(A)** Organs (heart, liver, kidney, lung, spleen, pancreas, and brain) were collected at D3 post *Leptospira* infection and stained for anti-LipL21. The scale bar corresponds to 20 µm for each organ except the brain where it corresponds to 50 µm. **(B)** Histology staining with Hematoxylin-Eosin (H&E) of brains of mice infected or not with *Leptospira* to detect inflammation. Open dots represent males and open squares represent females.

**Supplementary Figure 2.**
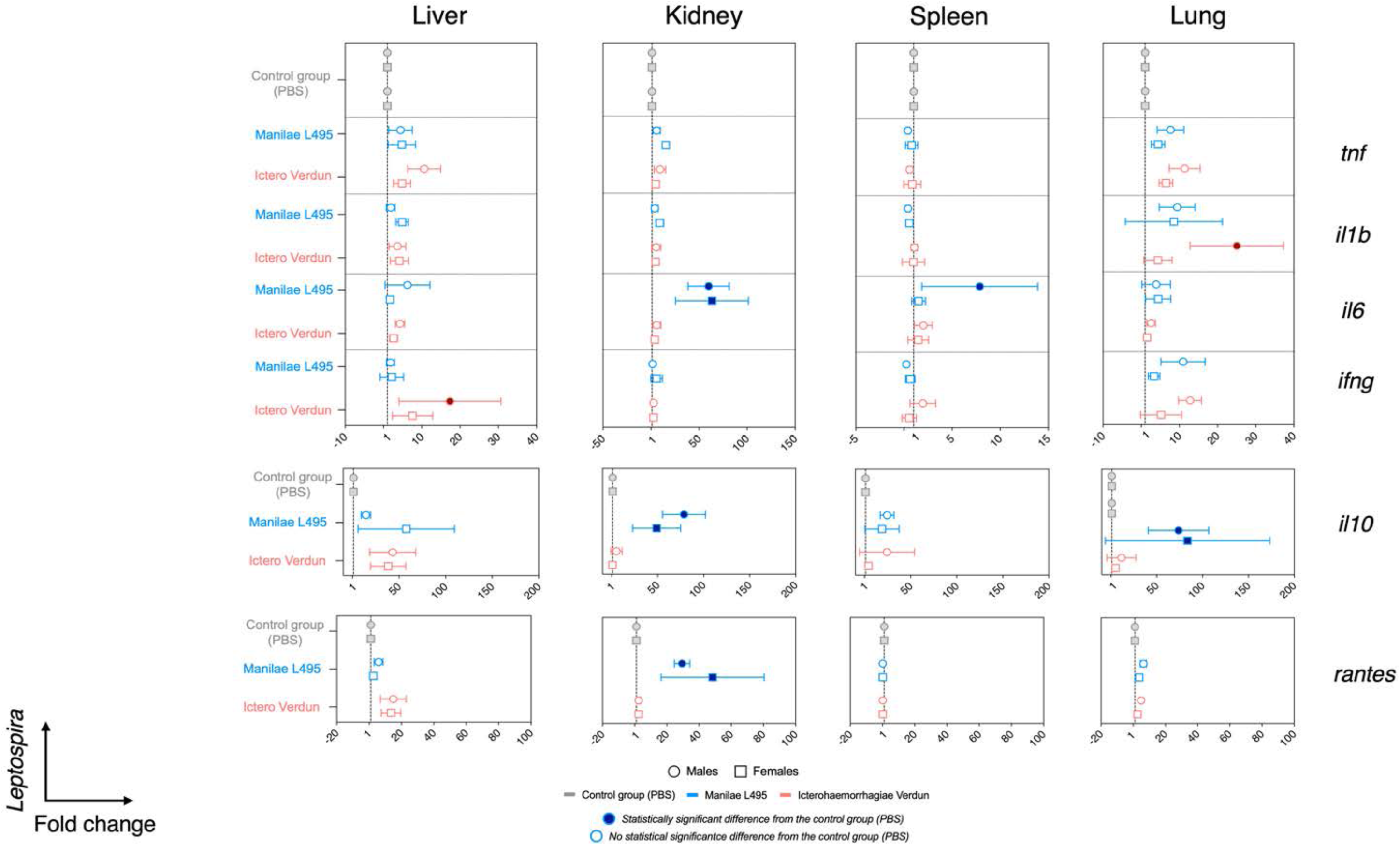
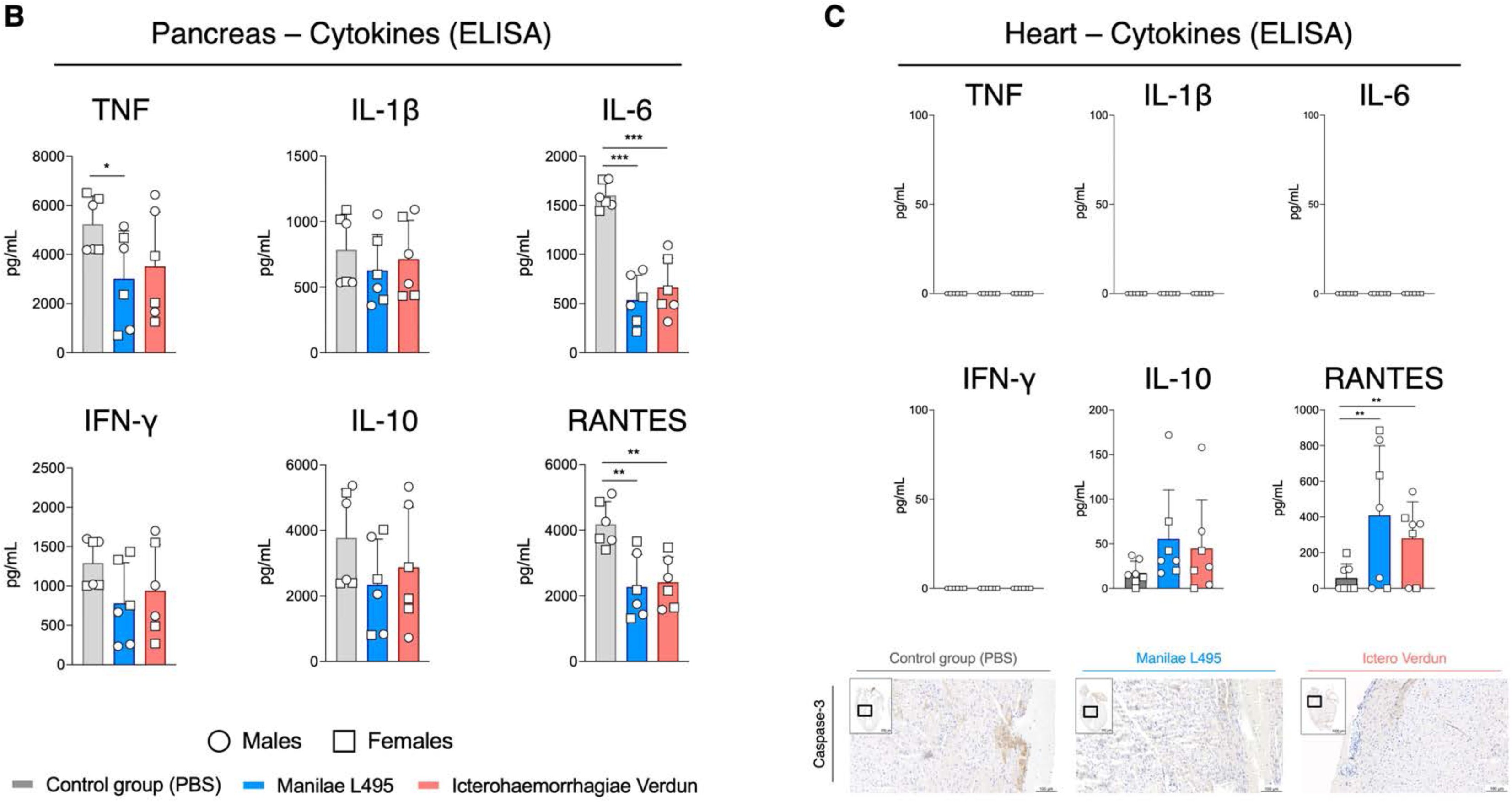
Quantification of cytokine mRNA in organs at the peak of the leptospiral infection and quantification of cytokine proteins during acute leptospirosis. **(A)** Organs (liver, kidney, spleen, lung) were collected at D3 post-*Leptospira* infection. Total RNA was isolated from organs, transcribed into cDNA and RT-qPCR was performed. The expression of pro-inflammatory *tnf*, *il1b*, *il6*, *ifng*, anti-inflammatory *il10* and chemokine *rantes* were determined with the 2^-ΔΔCt^ method. Each point of the forest plots represents the mean of n=5 per group. In dark bold colors are the statistically significant differences of the infected groups compared to the non-infected one. **(B)** Cytokine profile of pancreas at D3 p.i (n=6 per group). **(C)** *Upper part:* Cytokine profile of heart at D3 p.i (n=7 per group). *Lower part:* Immunohistochemical staining of the cardiac tissue at D3 post *Leptospira* infection with Caspase-3 to detect apoptosis. The pictures are representative of 3 independent experiments. Open dots represent males and open squares represent females.

**Supplementary Figure 3.**
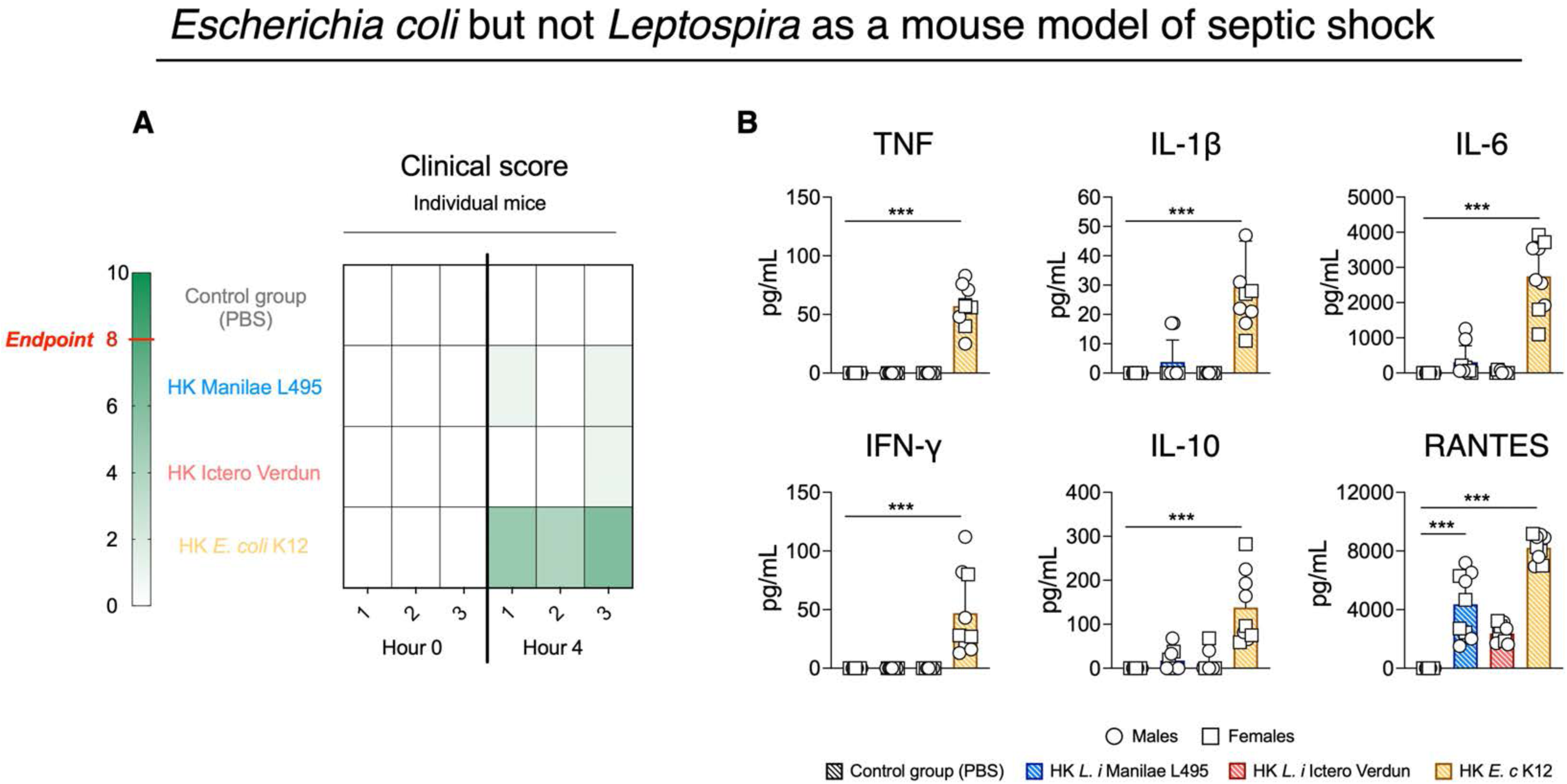
Mouse model of septic shock provoked by heat-inactivated *Escherichia coli, not Leptospira*. **(A)** Table showing the individual clinical scores in both male and female mice injected with heat-killed (HK) bacteria (*L. i* Manilae L495, *L. i* Icterohaemorrhagiae Verdun, *E. c* K12) or PBS (n=3 per group of 2 independent experiments), according to the hours post-infection, with color grading corresponding to severity. In dark green, the endpoint 8 corresponds to the humane ethical endpoint. **B)** Cytokine profile (Pro-inflammatory TNF, IL-1β, IL-6 and IFN-γ, anti-inflammatory IL-10 and chemokine CCL5/RANTES) determined by ELISA assay in plasma of mice collected 4 hours post-injection of HK bacteria. (n=9 per group of 2 independent experiments). Open dots represent males and open squares represent females.

**Supplementary Figure 4.**
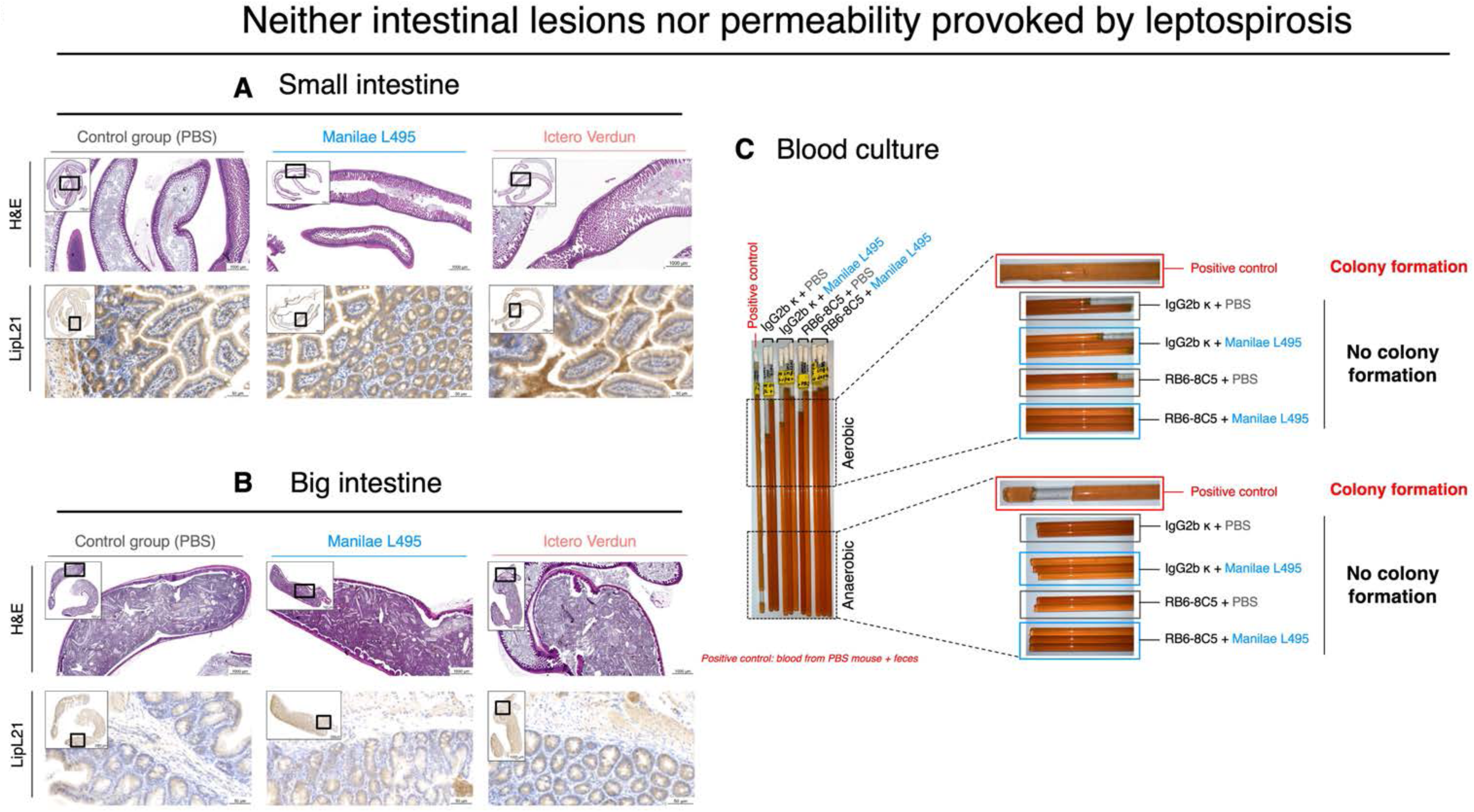
Leptospirosis does not provoke intestinal lesions nor permeability in mice. Histology staining of small intestine **(A)** and big intestine **(B)** of mice infected or not with *Leptospira* with Hematoxylin-Eosin (H&E) to detect inflammation, and immunohistochemistry of LipL21 to detect leptospires. The pictures are representative of 2 independent experiments. **(C)** Blood culture to detect intestinal permeability. Whole blood collected at D3 p.i from neutrophil-depleted or not mice and inoculated in BHI medium to detect the growth of aerobic/anaerobic bacteria. As positive control for colony formation, feces from naïve mice were mixed with the corresponding blood and then inoculated in BHI. The tubes remained at 37 °C for one week. The picture is representative of 2 independent experiments.

**Supplementary Figure 5.**
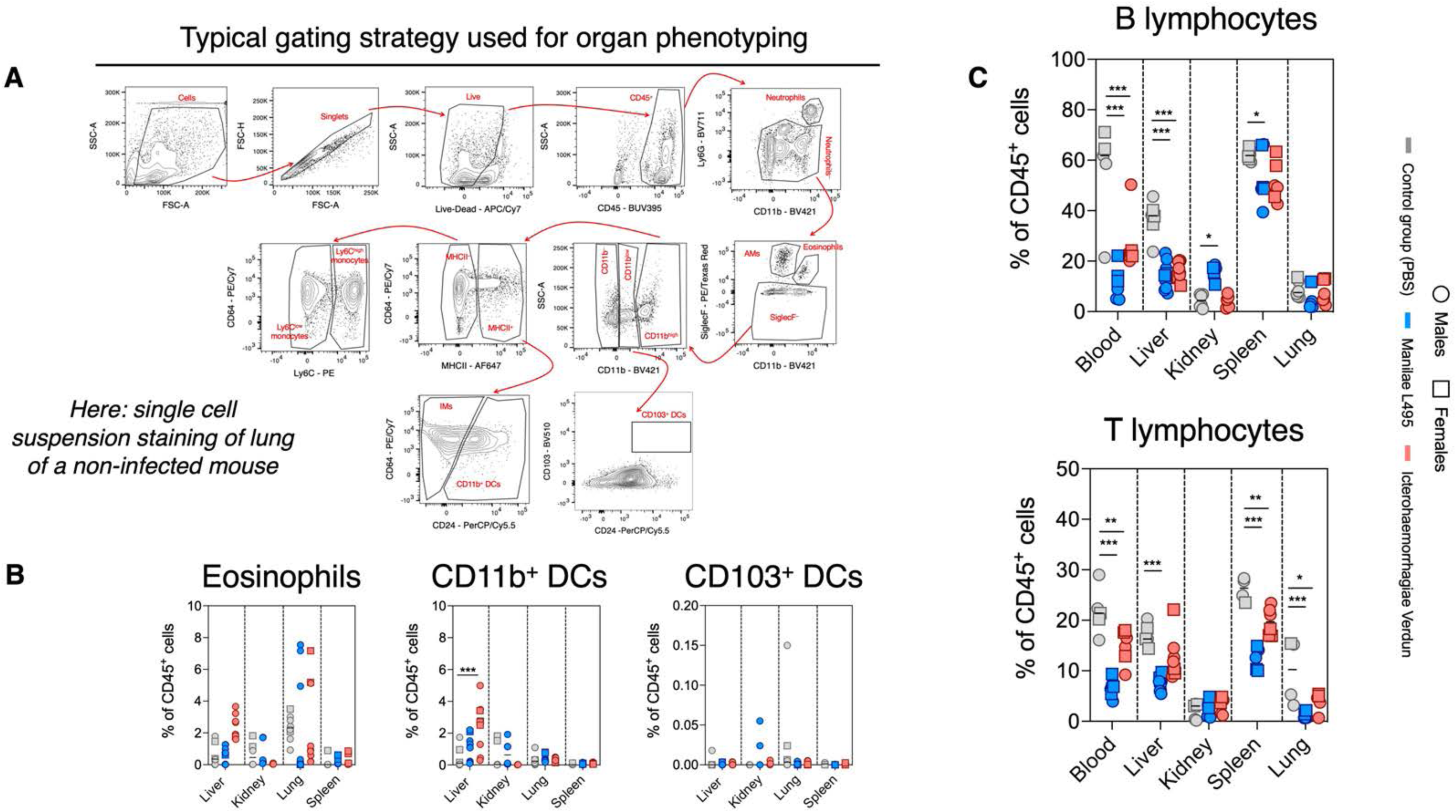
Cell populations upon phenotyping of organs. **(A)** Gating strategy followed for the phenotyping of liver, kidney, lung and spleen. In the current panel, the gating strategy is represented in the lung of a non-infected mouse. **(B)** Eosinophils, CD11b^+^ DCs and CD103^+^ DCs in liver, kidney, lung and spleen upon infection of mice with *Leptospira* (compilation of n=3 independent experiments). **(C)** Lymphocyte (B and T cells) in liver, kidney, lung and spleen upon infection of mice with *Leptospira* (compilation of n=3 independent experiments). Open dots represent males and open squares represent females.

**Supplementary Figure 6.**
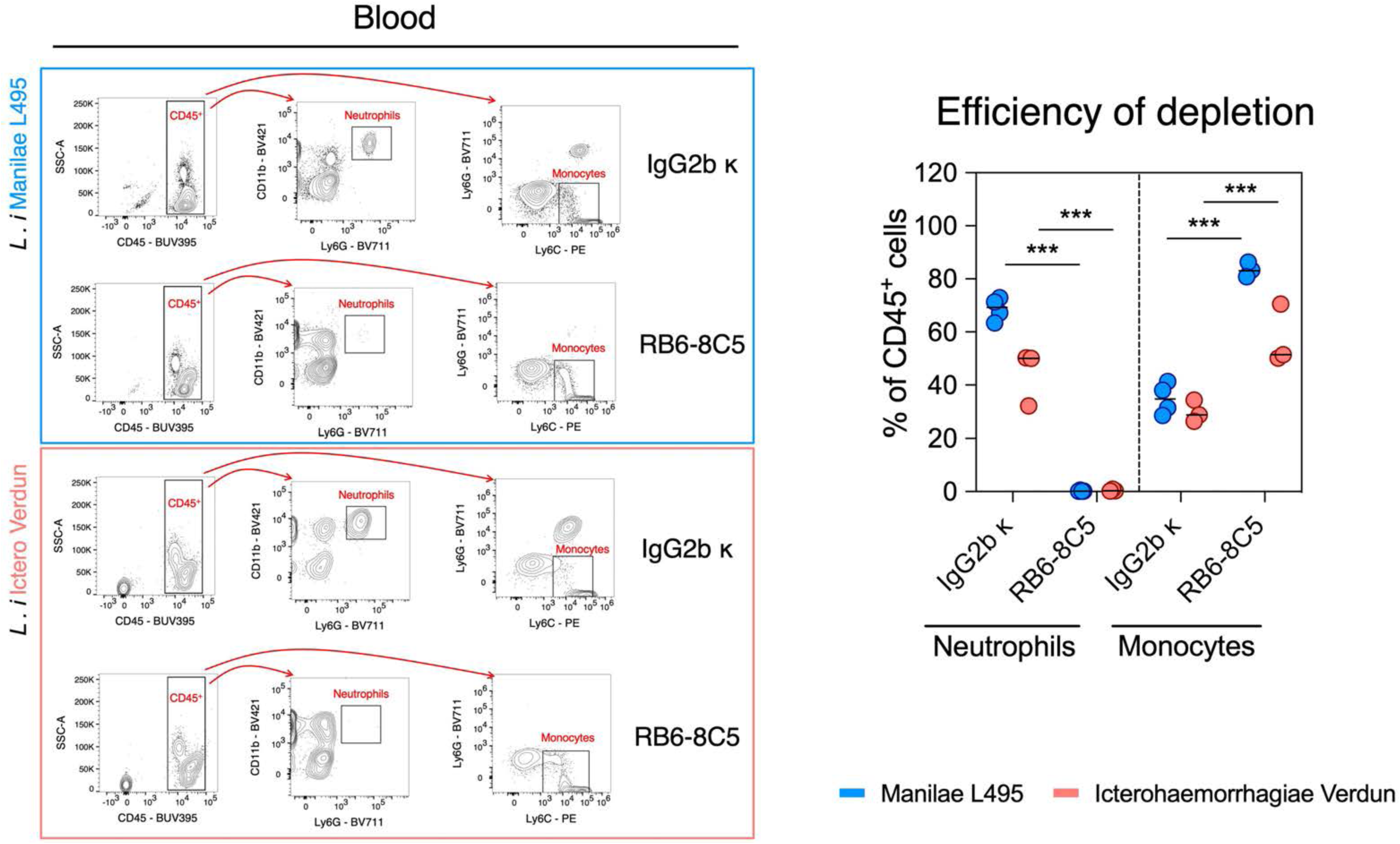
Efficiency of neutrophil depletion upon intravenal injection of RB6-8C5. **(A)** Gating strategy used to detect neutrophils and monocytes in whole blood. *Upper panel*: infection with *L. i* Manilae L495 and *lower panel*: infection with *L. i* Icterohaemorrhagiae Verdun. **(B)** Graph representing neutrophils and monocytes in the blood circulation of mice upon *Leptospira* infection in the presence (IgG2b κ) or absence (RB6-8C5) of neutrophils (representative graph of n=2 experiments).

## Notes

### Competing Interest Statement

The authors have declared no competing interest.

